# Power, positive predictive value, and sample size calculations for random field theory-based fMRI inference

**DOI:** 10.1101/613331

**Authors:** Dirk Ostwald, Sebastian Schneider, Rasmus Bruckner, Lilla Horvath

## Abstract

Recent discussions on the reproducibility of task-related functional magnetic resonance imaging (fMRI) studies have emphasized the importance of power and sample size calculations in fMRI study planning. In general, statistical power and sample size calculations are dependent on the statistical inference framework that is used to test hypotheses. Bibliometric analyses suggest that random field theory (RFT)-based voxel- and cluster-level fMRI inference are the most commonly used approaches for the statistical evaluation of task-related fMRI data. However, general power and sample size calculations for these inference approaches remain elusive. Based on the mathematical theory of RFT-based inference, we here develop power and positive predictive value (PPV) functions for voxel- and cluster-level inference in both uncorrected single test and corrected multiple testing scenarios. Moreover, we apply the theoretical results to evaluate the sample size necessary to achieve desired power and PPV levels based on an fMRI pilot study.

## 1. Introduction

A fundamental goal of task-related functional magnetic resonance imaging (fMRI) is to identify the cortical correlates of cognition. An approach routinely used to achieve this goal is mass-univariate null hypothesis significance testing in the framework of the general linear model (Friston et al., 1994; Poline and Brett, 2012; Cohen et al., 2017). In the recent debate on the reproducibility of research findings in the life sciences, the statistical practices of fMRI research have once again taken centre stage in the community discourse (e.g., Eklund et al., 2016; Mumford et al., 2016; Poldrack et al., 2017; Eklund et al., 2019; Flandin and Friston, 2019). Here, a particular emphasis has been on statistical power and its relation to typical sample sizes in fMRI group studies (Button et al., 2013; Guo et al., 2014; Szucs and Ioannidis, 2016; Cremers et al., 2017; Geuter et al., 2018; Turner et al., 2018). In task-related fMRI, statistical power is broadly defined as the probability of detecting cortical activation, if this activation is indeed present. In general, statistical power and, consequently, methods for computing the sample sizes necessary to achieve desired levels of power depend on both the statistical inference framework used and assumptions about the expected cortical activation.

A prominent statistical inference framework for null hypothesis significance testing in fMRI research is based on random field theory (RFT) (Worsley, 2007; Friston, 2007; Nichols, 2012; Ostwald et al., 2018). RFT-based fMRI inference is a parametric framework that allows for controlling the multiple testing problem inherent in the mass-univariate approach. Technically, this framework rests on analytical approximations to the exceedance probabilities of topological features of data roughness-adapted random field null models. RFT-based fMRI inference is implemented in the two major data analysis software packages used by the neuroimaging community, namely, Statistical Parametric Mapping (SPM) and the Functional Magnetic Resonance Imaging of the Brain (FMRIB) Software Library (FSL). It encompasses up to five forms of statistical testing: uncorrected and corrected voxel-level inference, uncorrected and corrected cluster-level inference, and set-level inference (Friston et al., 1996). With the exception of set-level inference, all forms are routinely reported in the functional neuroimaging literature. More specifically, bibliometric analyses suggest that RFT-based fMRI inference, especially corrected cluster-level inference, accounts for approximately 70% of published task-related human fMRI studies (Supplement S.1).

In light of the widespread use of RFT-based inference, previously proposed approaches for the calculation of power and sample sizes in fMRI research have a number of shortcomings. First and foremost, most previously proposed frameworks are not well aligned with the theory of RFT-based fMRI inference (e.g., Desmond and Glover, 2002; Mumford and Nichols, 2008; Durnez et al., 2016), rendering them non-applicable for the most commonly employed forms of fMRI inference. Second, the framework previously proposed by Hayasaka et al. (2007) and Joyce and Hayasaka (2012) that is aligned with the theory of RFT-based fMRI inference only addresses voxel-level and not cluster-level inference. Moreover, this framework does not address the variety of power types that arise in multiple testing scenarios and thus remains imprecise with respect to the interpretation of its ensuing power and sample size values. Third, all previous frameworks assume that under the alternative hypothesis, cortical activation is expressed either in a known region of interest or over the entire cortex. Notably, neither of these assumptions necessarily reflects common intuitions of neuroimaging researchers. Finally, no previous framework allows for the necessary sample sizes to be derived based on a desired positive predictive value (PPV), a novel statistical marker for the quality of empirical research that has risen to prominence over the last decade (Wacholder et al., 2004; Ioannidis, 2005; Heston and King, 2017; Colquhoun, 2019).

With the current work, we address these shortcomings and report on a novel framework for power, PPV, and sample size calculations in RFT-based fMRI inference. We first consider the framework’s theoretical foundations by briefly reviewing the notion of power in single test scenarios, the concepts of minimal and maximal power in multiple testing scenarios, the foundations of the PPV, and the notion of partial alternative hypotheses in Section 2: Theoretical foundations. We next develop the power and PPV functions for RFT-based fMRI inference on the basis of the standard fMRI group analysis probabilistic model in Section 3: Methods. In Section 4: Results, we then discuss the RFT-based power and PPV functions at both the voxel and cluster level in both the uncorrected single test and corrected multiple testing scenario and discuss their parametric dependencies. Finally, we apply the proposed framework in a prospective power analysis based on a pilot fMRI data set and evaluate the sample sizes necessary to obtain desired power and PPV levels. In Section 5: Discussion, we close with by considering some commonalities and differences between the proposed framework and previously proposed approaches and some potential avenues for future research. Throughout, we limit our scope to the evaluation of contrasts of first-level GLM parameter estimates (COPEs) at the group level using *T* -statistics, the approach most commonly used for group-level fMRI analyses. The technical foundations of our framework are detailed in Supplement S.2. All data and software used are available from https://osf.io/xjcg4/.

## 2. Theoretical foundations

### Power functions

In single test scenarios, such as testing for the activation of a single voxel, two types of errors can occur: the test may reJect the null hypothesis when it is in fact true, referred to as a Type I error, and the test may not reJect the null hypothesis when in fact the alternative hypothesis is true, referred to as a Type II error. From a frequentist perspective, Type I and Type II errors are associated with their probabilities of occurrence, denoted *α* and 1 − *β*, respectively, and commonly referred to as Type I and Type II error rates. The complementary probability of a Type II error, i.e., the probability rejecting the null hypothesis if the alternative hypothesis is true, is referred to as the power *β* of a test. A fundamental aim of test construction is to maintain low Type I and Type II error rates. To this end, a desired Type I error rate is usually selected first by defining a test significance level *α′*, ensuring a Type I error rate of at most *α′*. For many commonly used tests, the power at a fixed significance level *α′* can then be shown to be a function *β*(*n, d*) of an effect size measure *d* and the sample size *n*. An often recommended approach in research study design is calculating the necessary sample size *n* for which, under the assumption of a fixed effect size *d*, the power reaches a desirable level, such as *β*(*n, d*) = 0.8.

### Minimal and maximal power functions

In multiple testing scenarios, such as simultaneously testing for cortical activation over many voxels, a Type I or a Type II error may occur for each of the individual tests involved, inducing a variety of Type I and Type II error rates. For example, commonly considered Type I error rates in fMRI research are the *family-wise error rate* (FWER), defined as the probability of one or more false rejections of the null hypothesis, and the *false discovery rate* (FDR), defined as the expected proportion of Type I error among the rejected null hypotheses. Classically, the FWER has been the prime target for Type I error rate control in fMRI research. The prevalence of FWER control derives from the fact that the FWER can be efficiently controlled using maximum statistic-based procedures (e.g., Roy, 1953; Roy and Bose, 1953), which were at the centre of the early developments of RFT-based fMRI inference (Friston et al., 1991; Worsley et al., 1992; Friston et al., 1994). Maximum statistic-based multiple testing procedures allow the FWER to be controlled using a family-wise error significance level 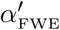 Just as the multiplicity of statistical tests in multiple testing scenarios induces a variety of Type I error rates, it also induces a variety of Type II error rates and hence power types. Power types commonly considered in multiple testing are *minimal power*, defined as the probability of one or more correct rejections of the null hypothesis, and *maximal power*, defined as the probability of correctly rejecting all false null hypotheses (e.g., Dudoit et al., 2003). When calculating the sample sizes necessary for desired power levels in Type I error rate-controlled multiple testing scenarios, it is hence essential to explicate the power type of interest. As RFT-based fMRI inference naturally lends itself to the evaluation of the minimal and maximal power functions *β*_*min*_(*n, d*) and *β*_*max*_(*n, d*), respectively, we focus on these power types in the current work.

### PPV functions

In recent discussions, studies with low power have been related to high probabilities of the claimed effects to be false positives (cf. Ioannidis, 2005; Button et al., 2013). This relationship is not inherent in classical frequentist test theory in which Type I and Type II error rates are conceived independently. Instead, the dependency of Type I error rates on Type II error rates, and hence power, arises in the context of a probabilistic model that assigns probabilities to the null hypothesis of being either true or false and the ensuing concept of a test’s PPV (Wacholder et al., 2004) (for an equivalent formulation in terms of false positive risk, see Colquhoun (e.g. 2017, 2019)). A test’s PPV, denoted here by ψ, is defined as the probability of the null hypothesis being false given that the test rejects the null hypothesis. As discussed in Supplement S.2, the PPV depends on both the Type I error rate and the *prior hypothesis parameter π* ∈ [0, 1], which represents the prior probability of the alternative hypothesis being true. For a constant Type I error rate and prior hypothesis parameter, the PPV is a function of the test’s power and, similar to power, a function ψ(*n, d*) of the effect and sample sizes. Moreover, in multiple testing scenarios, such PPV functions can be generalized to minimal and maximal PPV functions *ψ*_*min*_(*n, d*) and *ψ*_*max*_(*n, d*) by substitution of the respective minimal and maximal power functions. Similar to power functions, single test and multiple testing PPV functions allow finding the sample size *n* for which, at a given effect size *d*, the PPV function reaches a desirable level, such as *ψ*(*n, d*) = 0.8.

### Partial alternative hypothesis scenarios

Previous approaches to the evaluation of power in fMRI inference have typically relied on the assumption that the experimental effect of interest is expressed in a known cortical region of interest, i.e., single test scenarios, (e.g., Desmond and Glover, 2002; Mumford and Nichols, 2008), or in multiple testing scenarios, across the entire cortical volume (e.g., Hayasaka et al., 2007; Joyce and Hayasaka, 2012). While there are situations in which prospective power analyses are reasonable under these assumptions, we here suggest that the evaluation of necessary samples sizes may often be desired although neither the precise location of an expected activation nor the activation of the entire cortical sheet is reasonably assumed. To this end, we propose to parameterize the power, PPV, and sample size calculations in multiple testing scenarios with a *partial alternative hypothesis parameter λ* ∈ [0, 1], which describes the assumed proportion of activated brain volume. Intuitively, for example, *λ* = 0.1 corresponds to the assumption that 10% of the cortex is truly activated. Formally, *λ* corresponds to the continuous spatial generalization of the alternative hypotheses ratio of multiple testing scenarios, as discussed in Supplement S.2. Note that if *λ* = 0, the minimal and maximal power are necessarily identically zero, as there are no true activations. Equivalently, if *λ* = 1, the FWER is necessarily zero, as there are no null activations.

## 3. Methods

Based on the theoretical foundations discussed above, we next develop the power and PPV functions for RFT-based fMRI inference on the basis of the standard fMRI group analysis probabilistic model. For a comprehensive review of RFT-based fMRI inference from first principles and with a particular focus on its SPM implementation, please refer to Ostwald et al. (2018). For a comprehensive review of the underlying test theory, please refer to Supplement S.2.

### Probabilistic model

Standard fMRI group analysis in the framework of the GLM is based on a two-level summary statistics approach. At the first-level, participant-specific MRI time series are analysed using voxel-wise convolution-based GLMs. The resulting participant- and voxel-specific COPEs are the data used for the second-level, continuous-space, discrete-data point model of RFT-based fMRI inference,

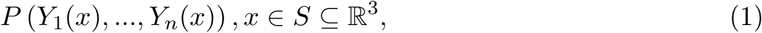

where *Y*_*i*_(*x*) denotes the random variable that models the COPE of the *i*th of *n* study participants at location *x* in the continuous three-dimensional search space *S*. In its structural form, the Joint distribution of these random variables is defined by

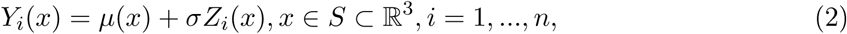

where *µ*(*x*) is an unknown value of a space-dependent parameter function *µ*: ℝ^3^ → ℝ, *σ* > 0 is an unknown standard deviation parameter, and *Z*_*i*_(*x*) is a *Z*-field modelling observation error. The *Z*_*i*_(*x*), *i* = 1, …, *n* are assumed to be independent and of identical smoothness. Observed COPE data sets are assumed to represent a lattice approximation to eq. (2) and can be represented by the discrete-space, discrete-data point model

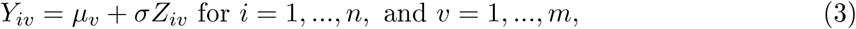

where *µ*_*v*_ := *µ*(*x*_*v*_) denotes the value of the parameter function *µ* at voxel location *x*_*v*_, and *Z*_*iv*_ := *Z*_*i*_(*x*_*v*_) denotes the *i*th *Z*-field random variable located at voxel location *x*_*v*_. In the following, we denote the family of random variables *Y*_*iv*_, *i* = 1, …, *n, v* = 1, …, *m* by *Y* := (*Y*_*iv*_)_*i*=1,…,*n,v*=1,…,*m*_, we summarize the values of the space-dependent effect size parameter function in a vector *µ* := (*µ*_1_, …, *µ*_*m*_)*^T^* ∈ ℝ^*m*^, and we denote the ensuing cardinality of the discretized second-level random field model and, equivalently, the dimensionality of an observed COPE data set, by *k* := *nm*.

### Statistics

RFT-based fMRI inference is based on a set of statistics that map *k*-dimensional COPE data sets onto lower-dimensional outcome spaces. Evaluating the probability of observed values of these statistics under the random field model of eq. (1) then allows for testing null hypotheses at desired levels of significance. To this end, RFT-based fMRI inference distinguishes single test scenarios, commonly referred to as *uncorrected inference*, based on *uncorrected p-values*, and multiple testing scenarios, commonly referred to as *corrected inference*, based on *corrected p-values*. Depending on the test scenario and the type of statistic, a specific form of inference ensues.

In the single test scenario and at the voxel level, the statistics of interest are the voxel height statistics

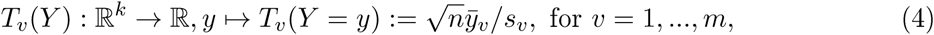

where 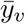 and *s*_*v*_ denote the sample mean and sample standard deviation of the *v*th voxel data, respectively. The voxel height statistics thus correspond to standard *T* -statistics and form so-called *statistical parametric maps* of *T* -statistics, sometimes denoted SPM{T}. Note that because under the *T* -statistic the Gaussian fields implied by (3) are projected onto a single *T* -field, the probabilities of statistics under the probabilistic model of eq. (1) are commonly expressed with respect to *T* -fields.

In the single test scenario and at the cluster level, the statistics correspond to the cluster extent statistics

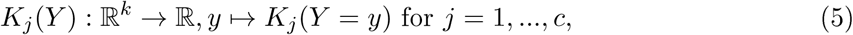

where *K*_*j*_ (*Y*) denotes the extent of the *j* th of *c* clusters within an excursion set defined by a CDT*u* ∈ℝ. The test statistics *K*_*j*_ (*Y*), *j* = 1, …, *c* subsume all data-analytical steps that project a COPE dataset onto the extents of clusters with in the excursion set of a statistical parametric map. These steps comprise but are not limited to thresholding a statistical parametric map at level *u*, evaluating the entailing clusters using a numerical connectivity scheme, and measuring the extent of the resulting clusters. Given the complexity of these computational subprocesses, closed-form expressions for the evaluation of *K*_*j*_ are not easily provided. Nevertheless, an approximation to the distribution of the test statistics *K*_*j*_ (*Y*), *j* = 1, …, *c* is routinely used in RFT-based FMRI inference, as will be discussed below.

In the multiple testing scenario and at the voxel level, the statistics of interest are the maximum and minimum of the voxel height statistics

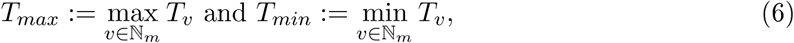

respectively. Similarly, in the multiple testing scenario and at the cluster level, the statistics of interest are the maximum and minimum of the cluster extent statistics

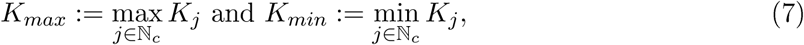

respectively. Consideration of the maximum statistics is warranted by their inherent property of enabling FWER control and the evaluation of minimal power in multiple testing scenarios. Consideration of the minimum statistics, in contrast, is warranted by their property of enabling the evaluation of maximum power in multiple testing scenarios. In the following, we detail the distributions of the statistics of eqs. (4)-(7) under the probabilistic model of eq. (1) that forms the core of RFT-based fMRI inference and the power evaluation framework proposed here. The distributions of the statistics will be provided in terms of exceedance probability functions (EPFs). EPFs are the probabilistic complements of cumulative probability functions and formulate the probability that a given statistic exceeds (rather than falls below, as in the case of cumulative probability functions) a given value. The use of EPFs is conventional in RFT-based fMRI inference and is useful in the contexts of false positive control and statistical power, both of which correspond to probabilities that statistics exceed critical values.

### EPFs of RFT-based fMRI inference statistics

The EPFs of the test statistics (4) - (7) are based on (1) the *T* -field’s search space resel volumes, (2) the *T* -field’s EC densities, and (3) three topological feature expectations. We discuss each of these in turn.

(1) The resel volumes

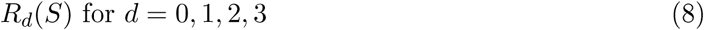

of a *T* −field’s search space *S* are the search space’s roughness-adjusted intrinsic volumes. In the SPM implementation of RFT-based fMRI inference, the resel volumes of a statistical parametric map are approximated by combining the values of the map’s intrinsic volumes with a standardized residuals-based roughness estimate using an algorithm originally proposed by Worsley et al. (1996).

(2) The EC densities of *T* -fields were originally derived as generalizations of the *T* -distribution by Worsley (1994). Based on work by Taylor et al. (2006), Hayasaka (2007) and Hayasaka et al. (2007) extended the EC densities to their non-central counterparts. The non-central *T* -field EC densities relevant for the current work are given by Hayasaka et al. (2007, p.729) as

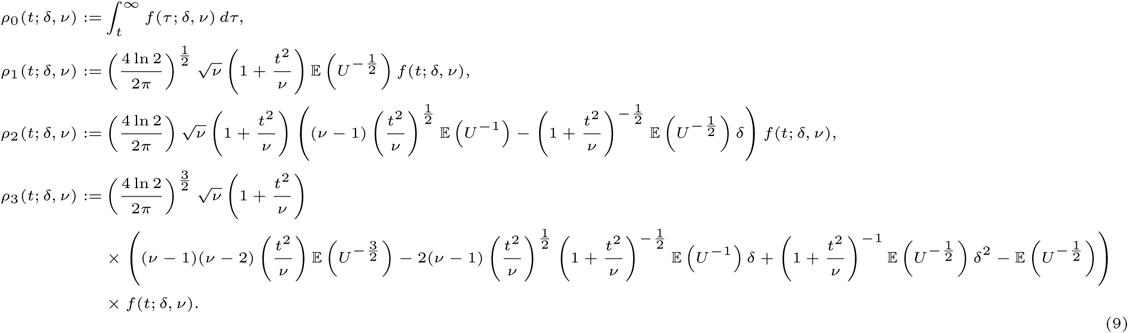

In eq. (9), *f* (*t*; *δ, ν*) denotes the probability density function of a non-central *T* random variable with non-centrality parameter *δ* ∈ ℝ and *ν* > 1 degrees of freedom, which is given by (e.g., Lehmann, 1986, p. 254, eq. (80))

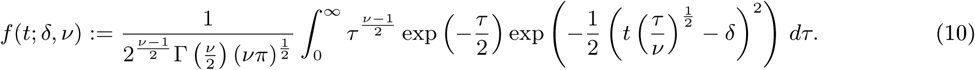

𝔼 (*U*^*p*^) denotes the expected value of the *p*th power of a non-central chi-squared random variable *U* with non-centrality parameter *ϕ* = *δ*^2^ and *µ* = *ν* + 1 degrees of freedom, which is given by (e.g., Johnson et al., 1995, p.449, eq. (29.32c))

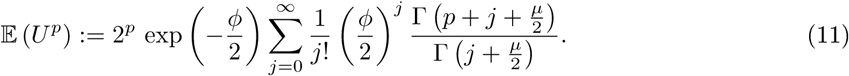

Note that for *δ* = 0, the non-central *T* -field EC densities (9) are identical to the (central) *T* - field EC densities as originally reported by Worsley et al. (1996) and Worsley et al. (1996). For the current work, the non-central *T* -field EC densities (9) are evaluated by the function r _ fun.m. This function computes *f* (*t*; *δ, ν*) using MATLAB’s nctpdf.m function, computes the integral of the zero-order non-central *T* -field EC density using Matlab’s nctcdf.m function, and approximates the series of eq. (11) by a numerically converging finite sum.

(3) Finally, the EPFs of the test statistics (4) - (7) are based on the following three topological feature expectations of *T* -fields: the expected volume of an excursion set, the expected number of clusters within an excursion set, and the expected volume of clusters within an excursion set. For the non-central *T* -field EC densities of eq. (9) with non-centrality parameter 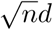 and *n* − 1 degrees of freedom, and for a CDT *u*, these expected values are given by

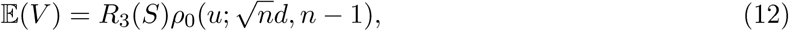

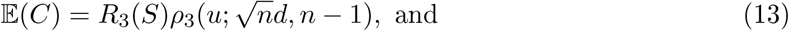

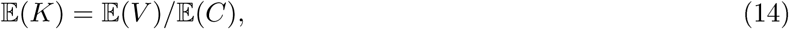

respectively.

With these preliminaries, the following EPFs for the statistics of eqs. (4) - (7) ensue:

○ The EPF of the voxel height statistics *T*_*v*_ follows from the standard theory of *T* -statistics. Moreover, because the zero-order non-central *T* -field EC density is identical to the cumulative density function of a non-central *T* -distribution, the EPF of the *T*_*v*_ for a non-central *T* –field with non-centrality parameter 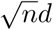 and *n* − 1 degrees of freedom takes the form

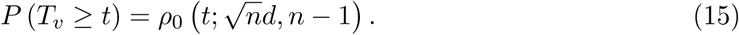 Note that for *d* = 0, the EPF of *T*_*v*_ equals the EPF of Student’s *T* -distribution with *n* − 1 degrees of freedom.
○ The EPF of the cluster extent test statistics *K*_*j*_ derives from an approximation for Gaussian random fields originally proposed by Friston et al. (1994). For a non-central *T* -field with non-centrality parameter 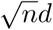 and *n* − 1 degrees of freedom, and for a CDT *u*, this approximation generalizes to

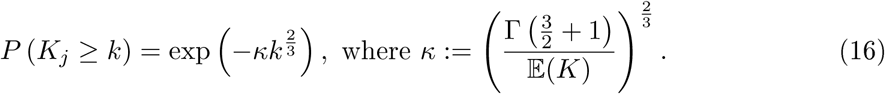
○ An approximation to the EPF of the maximum voxel height statistic *T*_*max*_ was originally proposed by Worsley et al. (1996) and was generalized to non-central *T* -fields by Hayasaka (2007). For a non-central *T* -field with non-centrality parameter 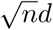 and *n* − 1 degrees of freedom, the approximation is given by

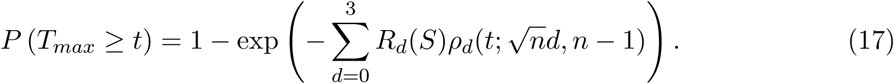 Similarly, as shown in Supplement S.3, an approximation to the EPF of the minimum voxel height statistic *T*_*min*_ can be given as

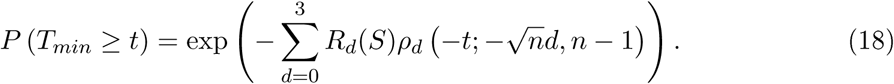
○ Finally, an approximation to the maximum cluster-level statistic *K*_*max*_ was proposed by Friston et al. (1994). Based on Hayasaka et al. (2007), this approximation can be generalized to non-central *T* -fields with non-centrality parameter 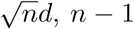,degrees of freedom, and a CDT *u* as

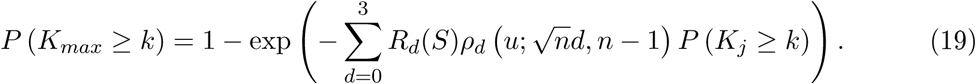 Similarly, as shown in Supplement S.3, an approximation to the EPF of the minimum cluster extent statistic *K*_*min*_ can be given as

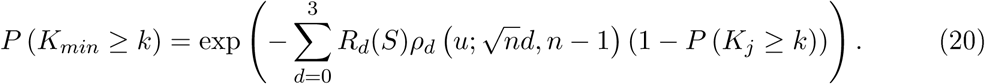

Test-relevant aspects of the EPFs in eqs. (15) - (20) are visualized in Supplement S.4. For all calculations, eqs. (15) - (20) are evaluated with the function q fun.m.

### Test hypotheses

The use of central *T* -field EC densities in the EPFs of fMRI inference test statistics reflects the intent to test the complete null hypothesis of zero activation throughout the entire search space *S*. Similarly, the use of non-central *T* -field densities in power calculations as proposed by Hayasaka et al. (2007) and Joyce and Hayasaka (2012) corresponds to the assumption of non-zero activation throughout the entire search space *S*. For our current work, we complement these boundary cases with the assumption of a parameterized partial alternative hypothesis scenario for power calculations. This scenario is based on the convex bipartition

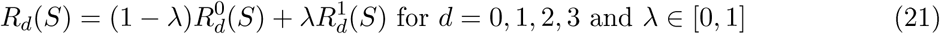

of the search space’s resel volumes *R*_*d*_(*S*), *d* = 0, 1, 2, 3 into resel volumes 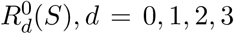 for which the null hypothesis of zero activation holds and resel volumes 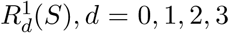 for which the alternative hypothesis of non-zero activation and with effect size parameter *δ* ≠ 0 holds. Note that for *λ* = 0, the partial alternative hypothesis scenario (21) corresponds to the complete null hypothesis of standard RFT-based fMRI inference, whereas for *λ* = 1, it corresponds to the complete alternative hypothesis scenario of Hayasaka et al. (2007) and Joyce and Hayasaka (2012). Intuitively, the value of *λ* thus corresponds to the proportion of the brain that is assumed to be activated for a given COPE. Formally, this proportion can be considered equivalent to the alternative hypothesis ratio in discrete multiple testing developed in Supplement S.2, eq. (S2.21). Specifically, for a partial alternative hypothesis parameter *λ* and a set of resel volumes *R*_*d*_, *d* = 0, 1, 2, 3, the expected Euler characteristic

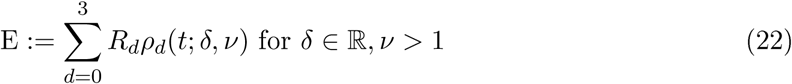

that combines resel volumes and EC densities in the EPFs of the RFT-based fMRI test statistics (4) - (7) takes the form

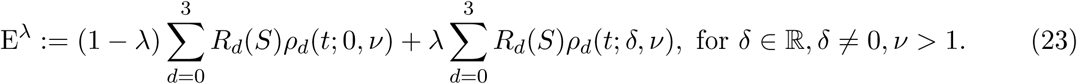

For eqs. (12) and (14), only the respective zero- and third-order terms are considered.

### Tests and power functions

With the test statistics and hypotheses in place, we next formalize the single test and multiple testing scenario for voxel- and cluster-level inference and document the power functions that result from the EPFs (15) - (20).

#### (1) Single test (uncorrected) voxel- and cluster-level inference

The aim of voxel-level inference in the single test scenario is to evaluate the null hypothesis of zero activation at the *v*th voxel location using the voxel height statistic *T*_*v*_ for the test

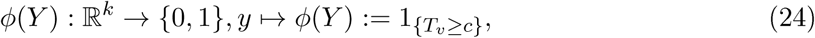

where 1_{·}_ denotes the indicator function and *c* denotes the test’s critical value. The Type I error rate of this test is controlled by choosing a critical value *t*_*α*_′ such that

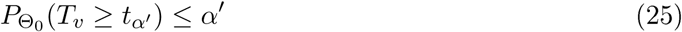

and the test obtains a significance level *α′*. With the EPF of *T*_*v*_, it then follows that the power function for voxel-level inference in the single test scenario is given by

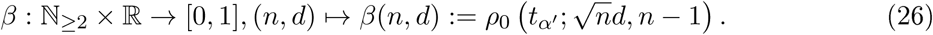

This power function corresponds to the standard power function for one-sample *T* -tests and is visualized in Figure 1A and Figure 1B. Note that the dependency of eq. (26) on the critical value *t*_*α ′*_ is commonly expressed indirectly in terms of the dependency of *t*_*α ′*_ on *α′* (cf. Figure 1B).

**Figure 1.**
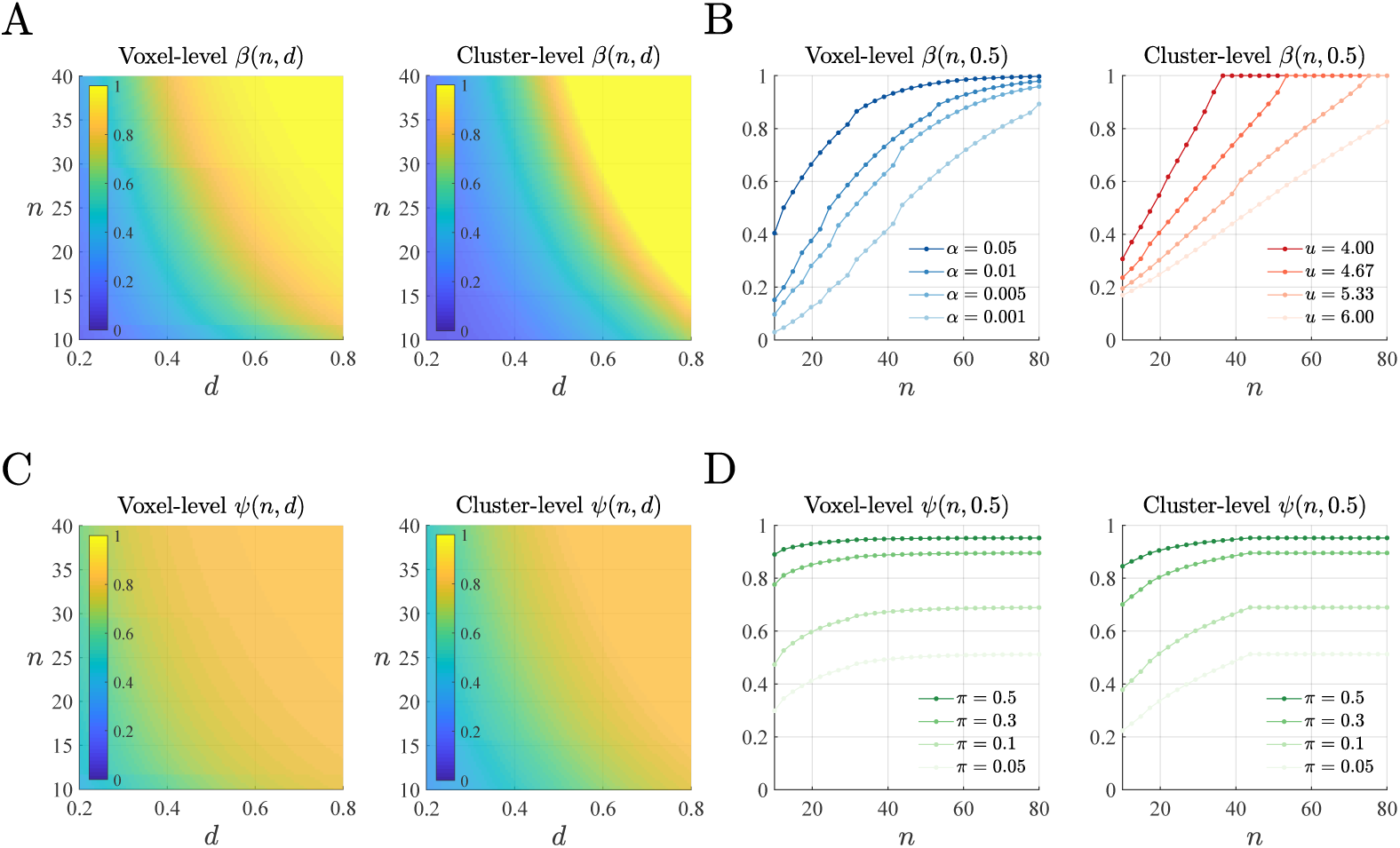
Power and PPV functions for voxel- and cluster-level inference in the uncorrected single test scenario. (A) Power functions for uncorrected voxel-level and cluster-level inference for a given sample size *n* and effect size *d*. For the cluster-level power function, a CDT parameter of *u* = 4.3 (*p* = 0.001 for *ν* = 9 degrees of freedom) was used. (B) Power dependency on the significance level *α*′ and the CDT value *u* for voxel- and cluster-level inference, respectively. (C) PPV functions for uncorrected voxel-level and cluster-level inference for a given effect size *d* and sample size *n* for the prior hypothesis parameter set to *π* = 0.2. (D) Prior parameter dependencies of the voxel- and cluster-level PPV functions for a fixed effect size of *d* = 0.5. Dots represent the evaluated sample sizes. For implementational details, please see rftp_figure_1.m.

The aim of cluster-level inference in the single test scenario is to evaluate the null hypothesis of zero activation over the extent of the *j*th cluster using the cluster extent statistic *K*_*j*_ for the test

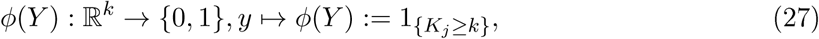

where *k* denotes the test’s critical value. The Type I error rate of this test is controlled by choosing a critical value *k*_*α ′*_ such that

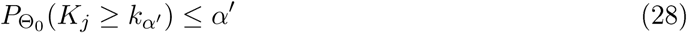

and the test obtains a significance level *α′*. With the EPF of *K*_*j*_, it then follows that the power function for the cluster-level inference in the single test scenario is given by

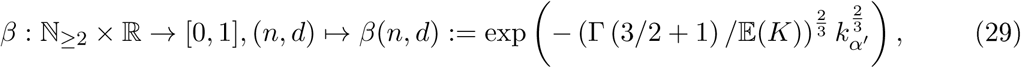

where

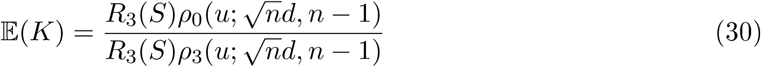

denotes the expected volume of a cluster in an excursion set at level *u*. This power function is visualized in Figure 1B.

#### (2) Multiple testing (corrected) voxel- and cluster-level inference

The aim of voxel-level inference in the multiple testing scenario is to evaluate the null hypothesis of zero activation at the *v*th voxel location while accounting for the multiplicity of tests over voxels using the multiple test

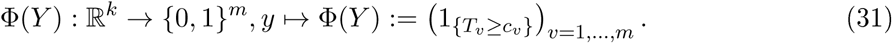

The FWER of this test is controlled based on the EPF of the maximum voxel height statistic *T*_*max*_ (17) by choosing a common critical value 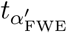 such that

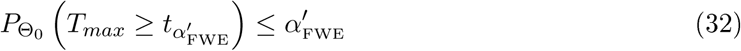

for a desired significance level 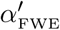 From the EPF of the maximum voxel height statistic (17), it then follows, that the minimal power function of voxel-level inference in the multiple testing scenario under the assumption of a partial alternative hypothesis with parameter *λ* is given by

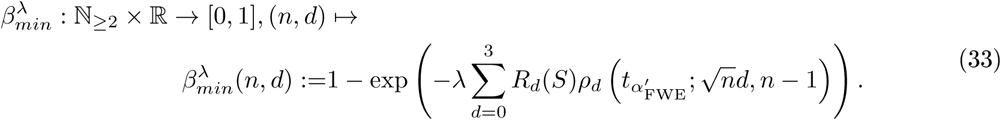

Similarly, from the EPF (18) of the minimum voxel height statistic *T*_*min*_, it follows that the maximal power function for voxel-level inference in the multiple testing scenario under the assumption of a partial alternative hypothesis parameter *λ* is given by

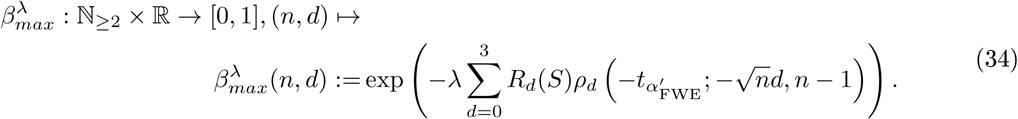

The ensuing minimal and maximal power functions for corrected voxel-level inference for *λ* = 0.1, 0.2, 0.3 are visualized in Figure 2A.

**Figure 2.**
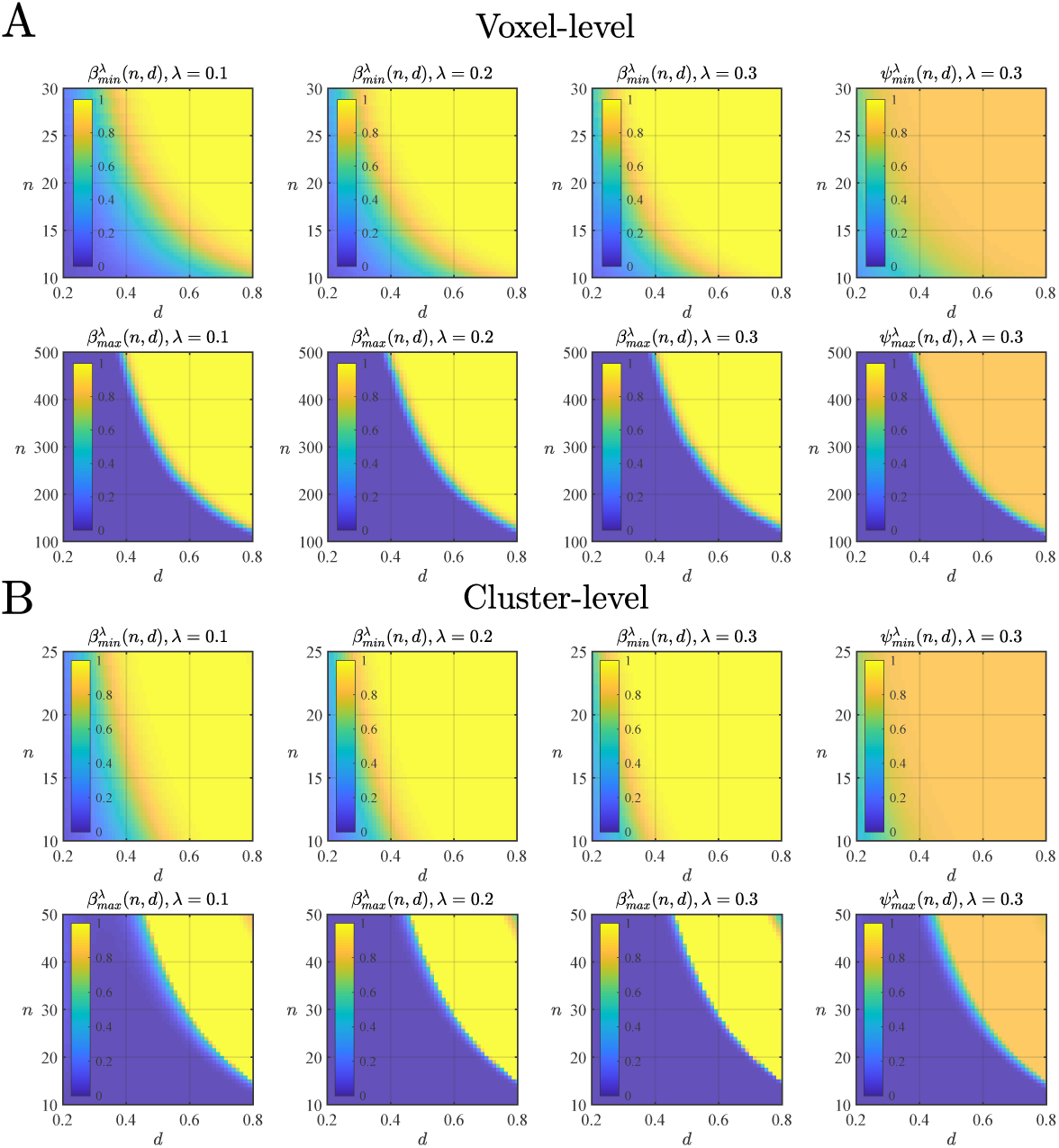
Minimal and maximal power and PPV functions for voxel- and cluster-level inference in the corrected multiple testing scenario. (A) Minimal and maximal power and PPV functions for corrected voxel-level inference for a given sample size *n*, effect size *d*, and partial alternative hypothesis parameter *λ* (first three columns). The fourth column depicts the corrected voxel-level minimal and maximal PPV functions for a prior hypothesis parameter of *π* = 0.2. (B) Minimal and maximal power and PPV functions for corrected cluster-level inference for a given sample size *n*, effect size *d*, and partial alternative hypothesis parameter *λ* (first three columns). The fourth column depicts the corrected cluster-level minimal and maximal PPV functions for a prior hypothesis parameter of *π* = 0.2. All cluster-level power functions were evaluated for a CDT of *u* = 4.3, and all voxel- and cluster-level power and PPV functions were evaluated for an exemplary resel volume set of *R*_0_ = 6, *R*_1_ = 33, *R*_2_ = 354, and *R*_3_ = 705. For further implementational details, please see rftp_figure_2.m.

Finally, the aim of cluster-level inference in the multiple testing scenario is to evaluate the null hypothesis of zero activation over the extent of the *j*th cluster location while accounting for the multiplicity of cluster tests using the multiple test

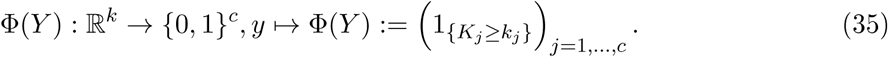

The FWER of this test is controlled based on the EPF of the maximum cluster extent statistic *K*_*max*_ (cf. (19)) by choosing a common critical value 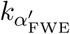 such that

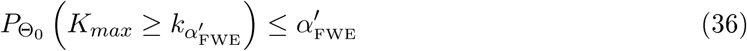

for a desired significance level 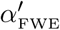. From the EPF (19) of *K*_*max*_, it then follows that the minimal power function of cluster-level inference in the multiple testing scenario under the assumption of a partial alternative hypothesis parameter *λ* is given by

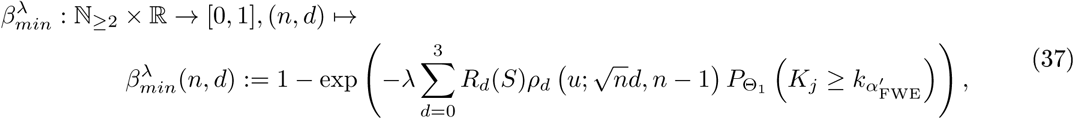

where 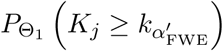 is evaluated according to (16) for resel volumes *λR*_*d*_, *d* = 0, 1, 2, 3. Similarly, from the EPF (20) of the minimum cluster extent statistic *K*_*min*_, it follows that the maximal power function for cluster-level inference in the multiple testing scenario under the assumption of a partial alternative hypothesis parameter *λ* is given by

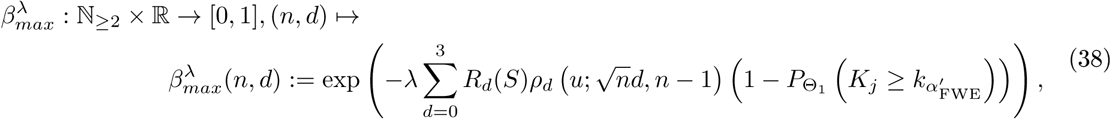

where 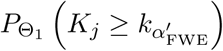 is evaluated as for 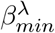 above. The ensuing power functions for *λ* = 0.1, 0.2, 0.3 are visualized in Figure 2B.

Note the differential manner by which the null hypothesis and alternative hypothesis resel volumes determine the minimal and maximal power functions in the multiple testing scenario: the null hypothesis resel volumes affect the determination of the critical values 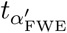 and 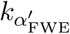 by means of the effective resel volumes (1−*λ*)*R*_*d*_(*S*) for some total resel volumes *R*_*d*_(*S*), *d* = 0, 1, 2, 3. In effect, the partial alternative hypothesis parameter *λ* here reduces the multiplicity of the multiple testing problem, as a control of the FWER is required (and possible) only over the resel volume subset (or clusters on this subset) for which the null hypothesis holds true. The alternative hypothesis resel volumes, in contrast, affect the evaluation of minimal and maximal power by means of the effective resel volumes *λR*_*d*_(*S*) for the same total resel volumes *R*_*d*_(*S*), *d* = 0, 1, 2, 3. If *λ* = 0, minimal and maximal power are identically zero, as there are no true activations. Equivalently, if *λ* = 1, there is no multiple testing problem and hence no FWER, as there are no non-activations. The power functions (26) - (38) are evaluated with the function p _ fun.m.

### PPV functions

As discussed in Supplement S.2, PPV functions for the five test scenarios of interest herein can be specified by means of the respective test’s (partial alternative hypothesis parameter-dependent) power function for sample size and effect size *β*(*n, d*), the test’s desired Type I error rate *α′*, and the prior hypothesis parameter *π* as

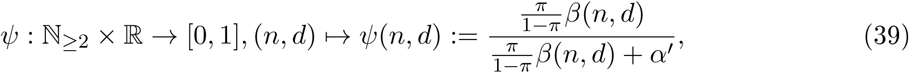

where the dependencies on *π* and *α′* are left implicit. The PPV functions depicted in Figure 1 - Figure 4 then follow directly by substituting the respective test power functions of eqs. (26),(33), (34), (29), (37), (38) in eq. (39)

**Figure 3.**
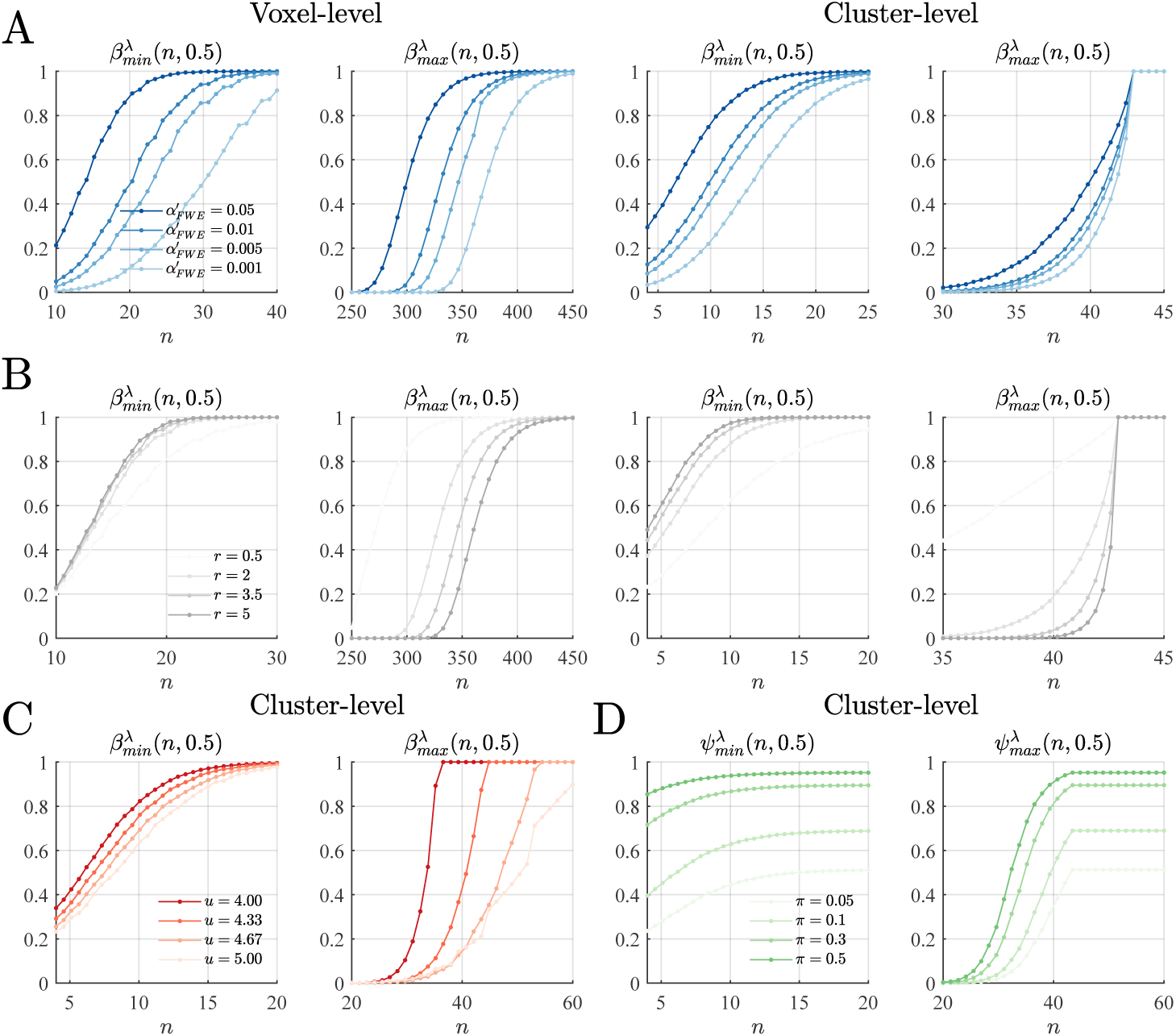
Parametric dependencies of minimal and maximal power and PPV functions for voxel- and cluster-level inference in the corrected multiple tesing scenario. Dots depict the evaluated sample sizes, and a medium effect size of *d* = 0.5 is considered for all plots. (**A**) Significance level dependencies of minimal and maximal power (*λ* = 0.1, *u* = 4.3). (**B**) Resel volume dependencies of minimal and maximal power (*λ* = 0.1, *u* = 4.3). *r* denotes the scalar multiple of the exemplary resel volume set of *R*_0_ = 6, *R*_1_ = 33, *R*_2_ = 354, and *R*_3_ = 705. (**C**) Minimal and maximal cluster-level power dependency on the CDT value *u*. (**D**) Prior hypothesis parameter dependencies of minimal and maximal PPV functions at the cluster level. For implementational details, please see rftp_figure_3.m.

**Figure 4.**
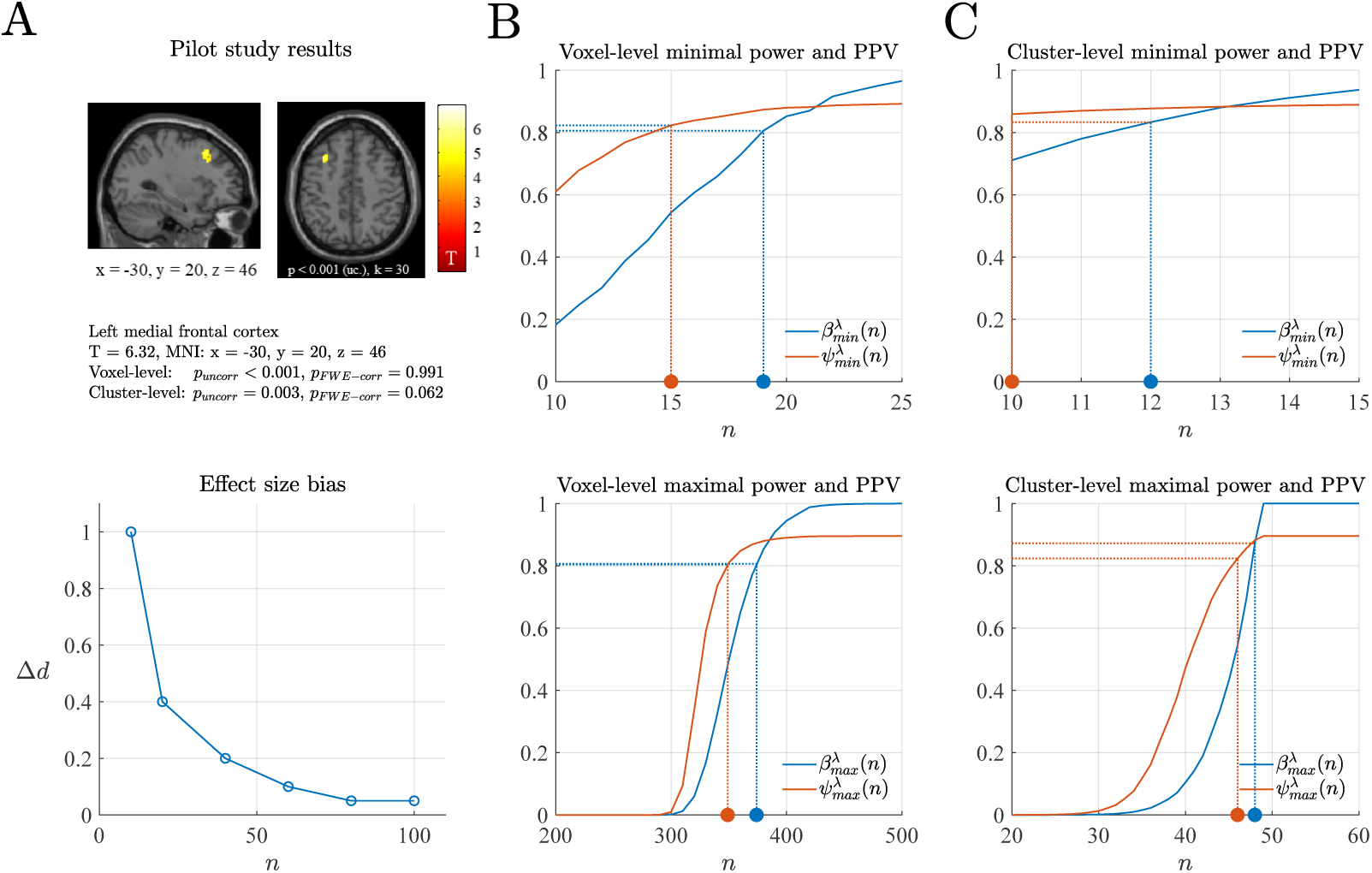
Exemplary application of the RFT-based power, PPV, and sample size calculation framework. (**A**) The upper panel depicts the results of a perceptual decision-making pilot study with *n* = 10 participants for contrasting perceptual choices based on high and low visual sensory evidence. The T-values from the identified cluster in the left medial frontal gyrus were averaged to obtain a raw effect size estimate, which was then adjusted based on the effect size bias estimates reported in Figure 7 of Geuter et al. (2018) and reproduced in the lower subpanel of panel (A). (**B**) Sample size calculations for voxel-level minimal and maximal power and PPV based on the effect size estimates of the pilot fMRI study. (**C**) Sample size calculations for cluster-level minimal and maximal power and PPV based on the effect size estimates of the pilot fMRI study. For implementational details, please see rftp_figure_4.m.

## 4. Results

### RFT-based fMRI inference power and PPV functions

In the following, we first discuss the power and PPV functions *β*(*n, d*) and ψ(*n, d*) for voxel- and cluster-level inference in single test scenarios for fixed significance levels *α′*. We then consider the power and PPV functions 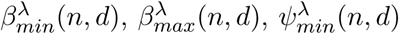 and 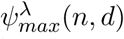 for voxel- and cluster-level inference for fixed family-wise error significance levels 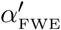 and for fixed partial alternative hypothesis parameters *λ*. Note that these functions form the essential prerequisites for calculating the sample sizes necessary to achieve desired levels of power or PPV.

### The single test scenario: uncorrected voxel- and cluster-level inference

Figure 1A depicts the power functions *β*(*n, d*) for voxel- and cluster-level inference in the uncorrected single test scenario at a significance level of *α′* = 0.05. For voxel-level inference and medium effect sizes of *d* = 0.4 to *d* = 0.6, sample sizes of *n* = 20 to *n* = 40 are required to achieve power levels of *β*(*n, d*) = 0.8. For cluster-level inference and similar effect sizes, slightly larger sample sizes of *n* = 25 to *n* = 40 are required to achieve similar power levels. Note that in contrast to voxel-level inference, cluster-level inference depends on the value of a cluster-defining threshold (CDT). For the cluster-level power function depicted in Figure 1A, the CDT was set to *u* = 4.3, corresponding to a *p*-value of 0.001 at *ν* = 9 degrees of freedom.

Naturally, varying the CDT impacts power: as shown in the right panel of Figure 1B, increasing the CDT at a constant sample size decreases power. This relationship is intuitive as, all else being equal, increasing the CDT will mask out an increasing number of voxels and hence reduce the chance of detecting a truly activated cluster. Similarly, and more fundamentally, the significance level impacts power for both voxel- and cluster-level inference: as depicted for voxel-level inference in the left panel of Figure 1B, decreasing the significance level decreases power. For all power curves shown in Figure 1B, the effect size was set to *d* = 0.5. For this medium effect size, a sample size of approximately *n* = 70 is required to achieve a power of *β*(*n, d*) = 0.8 at the uncorrected voxel-level significance level of *α′* = 0.001, which is sometimes used for inference in empirical studies. Notably, neither uncorrected voxel-level inference nor cluster-level inference is affected by the search space’s resel volumes that relate to the statistical map’s roughness: the RFT-based power function of the voxel-level height statistic (cf. eq. (26)) is identical to the power function of a one-sample *T* -test and is hence independent of the search space’s resel volumes per se. The power function of the cluster-extent statistic (cf. eq. (29)), however, is dependent on the expected cluster extent and hence potentially susceptible to variations in the statistical map’s roughness. However, as the third-order resel volume affects both the expected volume of clusters and the expected number of clusters, and for the evaluation of the expected cluster extent, RFT-based fMRI inference assumes the independence of these expectations (cf. eq. (12)), resel volume - and hence roughness - independence ensues.

Figure 1C depicts the PPV functions for voxel- and cluster-level inference in the uncorrected single test scenario as a function of effect size *d* and sample size *n* and for a prior hypothesis parameter of *π* = 0.2. Here, medium effect sizes similar to those of the power functions require sample sizes on the order of *n* = 10 to *n* = 30 and *n* = 15 to *n* = 35 to achieve PPV levels of ψ(*n, d*) = 0.8 for voxel- and cluster-level inference, respectively. From the definition of the PPV function ψ(*n, d*) as a monotonic transformation of a power function *β*(*n, d*) (cf. eq. (39)), it follows that the parameter dependencies of the voxel- and cluster-level power functions carry over to the respective PPV functions. Naturally, PPV functions are additionally strongly dependent on the value of the prior hypothesis parameter *π*: as shown in Figure 1D, low prior hypothesis parameter values result in much larger sample sizes necessary to achieve desired PPV levels, while higher prior hypothesis parameter values have the opposite effect.

### The multiple testing scenario: corrected voxel- and cluster-level inference

Figure 2A depicts maximal and minimal power and PPV functions for corrected voxel-level inference at a significance level of 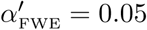. Specifically, the two leftmost panels of Figure 2A depict the minimal and maximal power functions 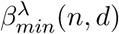 and 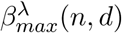 for corrected voxel-level inference and a partial alternative hypothesis parameter of *λ* = 0.1. Achieving a minimal power level of 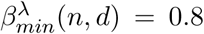. for a medium effect size of *d* = 0.5 requires sample sizes in the range of *n* = 15 to *n* = 30. To achieve similar levels of maximal power *β*_*max*_(*n, d*), the same effect size requires sample sizes of *n* = 200 to *n* = 500. As shown in the upper three panels of Figure 2A, increasing the partial alternative hypothesis parameter to *λ* = 0.2 and *λ* = 0.3 decreases sample sizes necessary to achieve a minimal power of 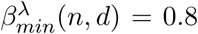. For maximal power, such a decrease is not observed. Intuitively, this relationship can be understood as follows: increasing the proportion of cortical activation increases the chances of detecting activation at a single cortical location (minimal power) but not of detecting activations at all locations (maximal power). Finally, for a prior hypothesis parameter of *π* = 0.2, PPV levels of 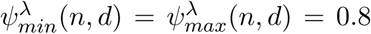. can be achieved with effect and sample sizes largely similar to those for minimal and maximal power, as depicted for *λ* = 0.3 in the rightmost column of Figure 2A.

Figure 2B depicts maximal and minimal power and PPV functions for corrected cluster-level inference at a significance level of 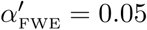. As for voxel-level inference, the leftmost panels of Figure 2B depict the minimal and maximal power functions for a partial alternative hypothesis parameter of *λ* = 0.1. Here, achieving a minimal power of 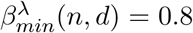 for a medium effect size of *d* = 0.5 requires sample sizes in the range of *n* = 10 to *n* = 20, while achieving a maximal power of 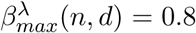 at the cluster level requires sample sizes of *n* = 30 to *n* = 50. As for corrected voxel-level inference, increasing the partial alternative hypothesis parameter to *λ* = 0.2 and *λ* = 0.3 decreases the necessary sample sizes for minimum power but not for maximum power. Finally, for a prior parameter of 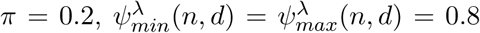 can also be achieved at the cluster level with effect and sample sizes largely similar to those for power (Figure 2B, rightmost column).

Naturally, the minimal and maximal power and PPV functions of corrected voxel- and cluster-level inference exhibit a number of additional parametric dependencies (Figure 3). First, as shown in Figure 3A, similar to the patterns observed for their uncorrected counterparts, the minimal and maximal power functions of corrected voxel- and cluster-level inference are affected by the desired significance level 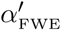, with lower values of 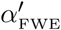 implying lower power. Second, and in contrast to the patterns observed for their uncorrected counterparts, the power functions in the corrected scenario are dependent on the data roughness, as expressed by a statistical map’s resel volumes. Figure 3B visualizes this influence as parameterized by a roughness parameter *r*, where for *r* = 1, the resel volumes are set as in Figure 2, while for *r* = 0.5 and *r* = 2 to *r* = 5, they are decreased or increased by the respective factor. Notably, for both voxel- and cluster-level inference, changes in the data roughness have opposite effects on minimal and maximal power: for minimal power, an increase in roughness *r* results in an increase of 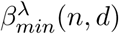, while for maximal power, an increase in roughness *r* results in a decrease of 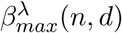. The effect of increased roughness on minimal power is familiar from the FWER-controlling features of the expected Euler characteristic (EC) (Adler, 1981; Worsley et al., 1996): the higher the roughness of the statistical field, the higher the probability for the maximum of the statistical field to exceed a given value, and hence the lower the statistical significance of an isolated peak. Because this relationship is a property of the maximum statistic *T*_*max*_, it is also evident in the case of minimal power. Intuitively, as the roughness of the statistical field can be viewed as a measure of the voxel height statistics’ spatial independence, detecting a single true alternative hypothesis is easier if it is not correlated with neighbouring height statistics. In contrast, maximal power increases with decreasing roughness and hence increasing smoothness. This association is intuitive: the smoother the statistical field is, the stronger the spatial covariation of the statistics. Thus, if a true alternative hypothesis is detected at one location, the other true alternative hypotheses are also likely to be detected (if, as in the current case, it is assumed that the area of activation corresponds to a contiguous set). As in the uncorrected cluster-level scenario, increasing the value of the CDT decreases power at a constant effect size for both minimal and maximal power (Figure 3C) because the probability of detecting one or all locations at which the alternative hypothesis is true decreases with the masking of an increasing number of voxels. Finally, the prior hypothesis parameter *π* also strongly affects PPV levels in the multiple testing scenario, as exemplified in Figure 3D for the cluster-level minimal and maximal PPV functions.

### Exemplary application

The power and PPV functions presented above imply the sample sizes necessary to achieve desired power and PPV levels over a broad range of possible effect sizes. To demonstrate the practical value of these functions, we finally consider their application in the concrete scenario of determining the sample size necessary to achieve power and PPV levels of 0.8 for a single effect size estimate. To this end, we re-analysed fMRI data from the first 10 participants in a previously reported perceptual decision-making study in which the amount of visual evidence for a presented stimulus to depict a face or a car was varied (Ostwald et al., 2012; Georgie et al., 2018). At the group level, contrasting fMRI activity levels between high and low visual evidence revealed a cluster of activity in the left medial frontal gyrus, as shown in the upper panel of Figure 4A (for further details about the experimental and data-analytical procedures, please see Supplement S.5). Our aim was to use the effect size estimate derived from this cluster to calculate the sample sizes necessary to achieve minimal and maximal power and PPV levels of 0.8 for corrected voxel- and cluster-level inference at a significance level of 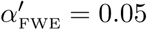, a partial alternative hypothesis parameter of *λ* = 0.1, and a prior hypothesis parameter of *π* = 0.2. To this end, we evaluated the average T-values of the cluster, yielding *T* = 4.65, which translates into an effect size estimate of 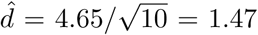. However, it is well known that effect size estimates resulting from the thresholding of mass-univariate statistical parametric maps exhibit biases (e.g., Vul et al., 2009; Poldrack et al., 2017). To correct our effect size estimate for this bias, we capitalized on recent results by Geuter et al. (2018), which are depicted in the lower panel of Figure 4A. Specifically, using task-related fMRI data from the Human Connectome ProJect 500 (Van Essen et al., 2013), Geuter et al. (2018) estimated the effect size bias exhibited by activations detected in random data subsets of 10 to 100 participants from the approximately 500 participants. As reported in Figure 7A of Geuter et al. (2018) and visualized in the lower panel of Figure 4A, this effect size bias is most severe for small data subsets and decreases with increasing data subset size. For a data subset of *n* = 10, the effect size bias amounts to approximately Δ*d* = 1. We thus used this empirically validated bias estimate to correct our effect size estimate to 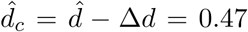. Using the power and PPV functions discussed in the previous section, and the sample size calculation algorithms Algorithm A2 and Algorithm A3 documented in Supplement S.6, we then obtained the following results: at the voxel level, sample sizes of *n* = 19 and *n* = 374 are required to achieve minimal and maximal power levels of 0.8, respectively (Figure 4B). At the cluster level, sample sizes of *n* = 12 and *n* = 48 are required to achieve minimal and maximal power levels of 0.8 (Figure 4C), respectively. For all testing scenarios considered and for the current parameter settings, slightly smaller sample sizes are required to achieve PPV levels of 0.8.

## 5. Discussion

In summary, we have developed power and PPV functions for RFT-based fMRI inference, which represents one of the mainstays of task-related fMRI data analysis. Further, we have demonstrated, how these functions can be used to determine the minimal sample sizes necessary to achieve desired power and PPV levels in study planning. Based on our example and its implementation in the MATLAB function rftp figure 4.m, interested users may readily adapt the procedures described herein for performing power, PPV, and sample size calculations in fMRI study planning. In the following, we briefly sketch the relation of the current framework to related approaches in the literature, discuss some potential avenues for future refinements of the approach, and close with some general remarks about statistical testing and power calculations in fMRI research.

The current framework can be thought of as a direct extension of the work by Hayasaka et al. (2007) and Joyce and Hayasaka (2012), generalizing the results presented therein to the cluster level and carefully distinguishing between uncorrected and corrected scenarios and the multiple power types thereby induced. As such, the current framework comprises region of interest-based approaches proposed by Desmond and Glover (2002) and Mumford and Nichols (2008) and implied in the discussions by Friston (2012) and Lindquist et al. (2013) as special cases. Specifically, in terms of its power function, a region of interest-based approach corresponds to uncorrected inference at the voxel level, i.e., a power evaluation for a one-sample *T* -test, with the difference that in typical region of interest-based approaches, voxel height statistics are spatially averaged over a set of voxels. Another power calculation framework that has recently been popularized is the approach of Durnez et al. (2016). This framework rests on a testing procedure that considers local maxima of voxel height statistics above a threshold. Under the model by Durnez et al. (2016), these local maxima are thought to be the outcome of a mixture distribution, comprising realizations of a null hypothesis exponential distribution and an alternative hypothesis Gaussian distribution. While the test procedure itself is not explicitly described, the apparent idea is to reJect the null hypothesis of no activation at the location of the local maximum based on a set of arbitrary selected critical values (Durnez et al., 2016, Section 3.3). Based on parameter estimates for the alternative hypothesis mixture component and the selected critical value, Durnez et al. (2016) calculate power and sample sizes. While an interesting approach in its own right, the method by Durnez et al. (2016) relates to statistical models and testing procedures that are specific to the power calculation approach by Durnez et al. (2016) and that are not routinely used in fMRI data analysis.

The current work implies some potential avenues for further research with the aim of improving power, PPV, and sample size calculations for fMRI inference. First, RFT-based fMRI inference itself may be further refined, thus entailing an optimization of the power and PPV framework discussed herein. For example, the approximations to the cluster-level test statistic distributions remain to be based on the Gaussian random field approximations by Friston et al. (1994), while newer results for *T* - and *F* -fields are available (e.g., Cao, 1999). Similarly, the notion of resel volumes has been largely superseded by the concept of Lipschitz-Killing curvatures (e.g., Taylor and Worsley, 2007), a theoretical development that has yet to be considered in standard discussions of RFT-based fMRI inference. Second, it has been observed previously as well as by us that some of the power functions of the RFT-based inference framework can behave non-monotonically outside of practically relevant parameter regimes (Hayasaka et al., 2007). Therefore, it may be desirable to further pursue mathematical analysis of the RFT-based exceedance probability function approximations and to study their analytic behaviour across parameter regimes. Finally, with respect to the PPV, it may be desirable to diminish the degree of subjectivity involved in selecting the prior hypothesis parameter. Potential avenues with which to achieve this goal include basing PPV calculations on empirical priors estimated from fMRI pilot data and considering the PPV in the more general setting of the false positive risk (e.g., Colquhoun, 2017, 2019).

As emphasized throughout, statistical power and PPVs are rooted in statistical testing, i.e., the dichotomization of the uncertainty-imbued results of statistical inference. As such, statistical testing, power and PPV calculations, as well as deriving the sample sizes necessary to achieve desired power and PPV levels, always generate simplified answers to complex scientific questions (e.g., Wasserstein et al., 2019). Such simplified answers may not always be desired in a scientific context, as indicated by recent initiatives to share unthresholded statistical parametric maps (Gorgolewski et al., 2015). Stated differently, while many researchers have argued that abandoning statistical testing based on arbitrary significance thresholds may be a promising avenue for improving scientific inference, few have argued that the entailing abandonment of power analyses may have similar effects. While we share the hope that the fMRI community will abandon statistical testing in the long run, we here have provided power, PPV, and sample calculations applicable to the widely used RFT-based fMRI inference procedures that can be adopted in the meantime.

## Author contributions

D.O. designed and performed the research and analysed the data. D.O. wrote the paper with input from S.S., R.B., and L.H.

## Acknowledgements

We thank Hauke Heekeren for comments on an earlier version of this manuscript.

## Data availability

All data used are available at https://osf.io/xjcg4/.

## Software availability

All software used is available at https://osf.io/xjcg4/.

## Supplementary Material

### S.1. Bibliometrics

Over the past seven years, at least four studies have used bibliometric methods and one study has used survey methods to assess the use of data analysis software packages and statistical testing procedures in the functional neuroimaging literature (Carp, 2012; Woo et al., 2014; Poldrack et al., 2017; Borghi and Van Gulick, 2018; Yeung, 2018). The most recent and most comprehensive account is provided by Yeung (2018). In Table S.1, we summarize the reported use of the SPM and FSL software packages for data analysis, the use of RFT-based methods for multiple testing control, and the relative prevalences of corrected voxel- and cluster-level inference. Note that because RFT-based inference is the default option in SPM, it is likely that the choice of the SPM software often implies the use of RFT-based inference, even if this is not explicitly stated in the primary research study nor the meta-research studies cited here. Also note that, as reported in Poldrack et al. (2017, p.123), up to a third of published fMRI studies continue to fail identifying the method used for multiple testing control, a fact which has prompted initiatives such as the COBIDAS report to improve reporting standards in the fMRI literature (Nichols et al., 2017).

**Table S1.**
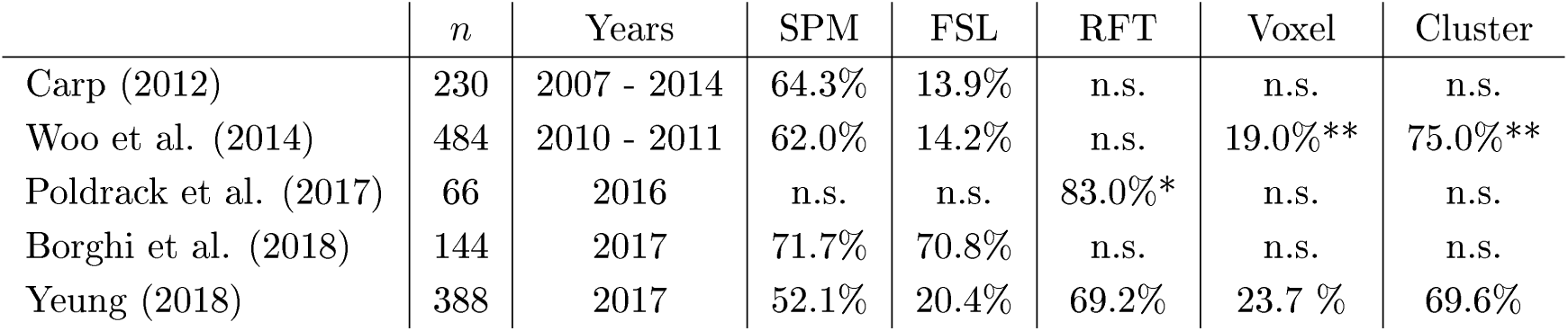
Bibliometric and survey data on the use of software packages and statistical inference procedures in the fMRI literature. All cited meta-research studies focus on human fMRI and assessed data analysis methods in *n* = 66 to *n* = 484 primary research studies published between 2007 and 2017. The study by Borghi and Van Gulick (2018) is based on survey data, all other studies used bibliometric methods. The study by Poldrack et al. (2017) assessed the most recently published articles as of May 2016, all other bibliometric studies specify the exact time range of the evaluated research reports. The prevalence of SPM and FSL use ranges between 52.1% and 71.7% and 13.9% and 70.8%, respectively. The use of RFT-based fMRI inference is explicitly mentioned in two of the meta-research studies and accounts for approximately 75% of the reported multiple testing control methods. Corrected cluster-level inference dominates corrected voxel-level inference by approximately 70% to 20% (*n*: number of studies or survey participants included, Years: time range of the assessed literature or active use, SPM: Statistical parametric mapping software, FSL: FMRIB software library, RFT: random field theory use for multiple testing control, Voxel: corrected voxel-level inference, Cluster: corrected cluster-level inference, n.s.: not, or insufficiently, specified, *: estimated based on the verbose description on p.122 of Poldrack et al. (2017), **: based on *n* = 814 primary research studies).

| | $n$ | Years | SPM | FSL | RFT | Voxel | Cluster |
| --- | --- | --- | --- | --- | --- | --- | --- |
| Carp (2012) | 230 | 2007 - 2014 | 64.3% | 13.9% | n.s. | n.s. | n.s. |
| Woo et al. (2014) | 484 | 2010 - 2011 | 62.0% | 14.2% | n.s. | 19.0%** | 75.0%** |
| Poldrack et al. (2017) | 66 | 2016 | n.s. | n.s. | 83.0%* | n.s. | n.s. |
| Borghi et al. (2018) | 144 | 2017 | 71.7% | 70.8% | n.s. | n.s. | n.s. |
| Yeung (2018) | 388 | 2017 | 52.1% | 20.4% | 69.2% | 23.7 % | 69.6% |

### S.2. Test theory

In this Section we review the formal foundations of test theory. We first develop the single hypothesis test scenario and its associated error rates and power function. We then consider the multiple testing scenario with a particular emphasis on the notions of partial alternative hypothesis scenarios as well as minimal and maximal power functions. In a third step, we discuss the probabilistic foundations of the positive predictive value. We close our review by discussing a single-observation z-test in the context of the single test and the multiple testing scenario.

#### S.2.1. The single test scenario

##### PROBABILISTIC MODEL

To introduce the notion of a single test, we consider a parametric probabilistic model *P*_*θ*_(*Y*) that describes the probability distribution of a random entity (i.e., a random variable or a random vector) *Y* and that is governed by a parameter *θ* ∈ Θ. The random entity *Y* models *data* and is assumed to take on values *y* ∈ ℝ^*n*^, *n* ≥ 1. Note that we do not consider the parameter *θ* to be a random entity and thus develop the following theory against the background of the classical frequentist scenario.

##### TEST HYPOTHESES

In test scenarios, the parameter space Θ is partitioned into two disjoint subsets, denoted by Θ_0_ and Θ_1_, such that Θ = Θ_0_ *∪* Θ_1_ and Θ_0_ *∩* Θ_1_ = ∅. A test *hypothesis is* a statement about the parameter governing *P*_*θ*_(*Y*) in relation to these parameter space subsets. Specifically, the statement

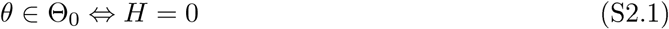

is referred to as the *null hypothesis* and the statement

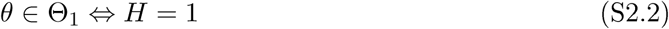

is referred to as the *alternative hypothesis*. Note that we are concerned with the Neyman-Pearson hypothesis testing framework and thus assume that null and alternative hypotheses always exist in an explicitly defined manner. A number of things are noteworthy. First, a statistical hypothesis is a statement about the parameter of a probabilistic model. In the following, we will use the subscript notations 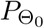 and 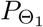 to indicate that the parameter *θ* of the probabilistic model *P*_*θ*_ is an element of Θ_0_ or Θ_1_, respectively. Second, the term null hypothesis is not necessarily the statement that some parameter assumes the value zero, even if this is often the case in practice. Rather, the null hypothesis in a statistical testing problem is the statement about the parameter one is willing to nullify, i.e., reject. Finally, the expressions *H* = 0 and *H* = 1 are not conceived as realizations of a random variable and hence hypothesis-conditional probability statements are not meaningful. The statements *H* = 0 and *H* = 1 are merely equivalent expressions for *θ* ∈ Θ_0_ and *θ* ∈ Θ_1_, respectively: *H* = 0 refers to the true, but unknown, state of the world that the null hypothesis is true and the alternative hypothesis is false (*θ* ∈ Θ_0_), and *H* = 1 refers to the true, but unknown, state of the world that the alternative hypothesis is true and the null hypothesis is false (*θ* ∈ Θ_1_). In general, hypotheses can be classified as *simple or composite*. A *simple hypothesis* refers to a subset of parameter space which contains a single element, for example Θ_0_ := *{θ*_0_*}*. A *composite hypothesis* refers to a subset of parameter space which contains more than one element, for example Θ_0_ := ℝ_*≤*0_. The commonly encountered *null hypothesis* Θ_0_ = *{*0*}*, also referred to as nil hypothesis, is an example for a simple hypothesis.

##### TESTS

Given the test hypotheses scenario introduced above, a test is defined as a mapping from the data outcome space to the set {0, 1}, formally

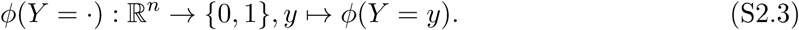

Here, the test value *ϕ*(*Y* = *y*) = 0 represents the act of not rejecting null hypothesis, while the test value *ϕ*(*Y* = *y*) = 1 represents the act of rejecting the null hypothesis. Rejecting the null hypothesis is equivalent to accepting the alternative hypothesis, and accepting the null hypothesis is equivalent to rejecting the alternative hypothesis. In the following and in the main text, we suppress the notational dependence of *ϕ*(*Y* = *·*) on *y* and write *ϕ*(*Y*) instead. Because *Y* is a random entity, the expression *ϕ*(*Y*) is also a random entity. All tests *ϕ*(*Y*) considered in the current study involve the composition of a *test statistic*

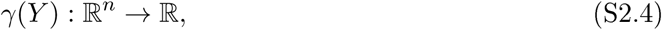

where ℝ models the test statistic’s outcome space, and a subsequent *decision rule*

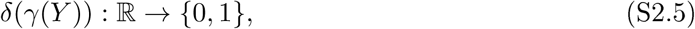

such that the test can be written as

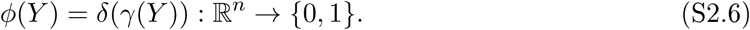

Note that, as for the test, we suppress the dependencies of *γ*(*Y*) and *δ*(*γ*(*Y*)) on *y* ∈ ℝ^*n*^, such that both *γ*(*Y*) and *δ*(*γ*(*Y*)) should be read as random entities. The subset of the test statistic’s outcome space for which the test assumes the value 1 is referred to as the rejection region of the test. Formally, the *rejection region* is defined as

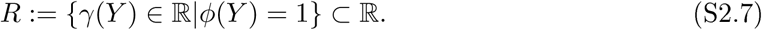

The random events *ϕ*(*Y*) = 1 and *γ*(*Y*) ∈ *R* are thus equivalent and associated with the same probability under *P*_*θ*_(*Y*). In a concrete test scenario, it is hence usually the probability distribution of the test statistic that is of principal concern for assessing the test’s outcome behaviour. Finally, all test decision rules considered in the context of the current study are based on the test statistic exceeding a *critical value u* ∈ ℝ. By means of the indicator function, the tests considered here can thus be written

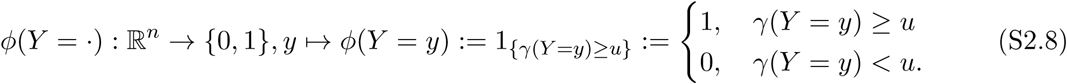

Note that (S2.8) describes the situation of *one-sided* tests. The one-sided one-sample *T* -test is a familiar example of the general test structure described by expression (S2.8): using the sample mean and sample standard deviation, a realization of the random entity *Y* is first transformed into the value of the t-statistic, whose size is then compared to a critical value in order to decide for rejecting the null hypothesis or not.

##### TESTS ERROR PROBABILITIES

When conducting a hypothesis test as Just described, two kinds of errors can occur. First, the null hypothesis can be rejected (*ϕ*(*Y*) = 1), when it is in fact true (*θ* ∈ Θ_0_). This error is referred to as the *Type I error*. Second, the null hypothesis may not be rejected (*ϕ*(*Y*) = 0), when it is in fact false (*θ* ∈ Θ_1_). The latter error is known as the *Type II error*. The probabilities of Type I and Type II errors under a given probabilistic model are central to the quality of a test: the probability of a Type I error is called the size of the test and is commonly denoted by *α* ∈ [0, 1]. It is defined as

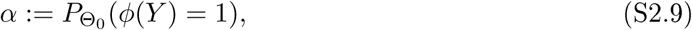

and also routinely referred to as the *Type I error* rate of the test. Its complementary probability,

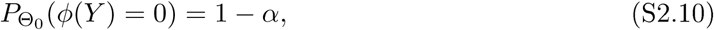

is known as the *specificity* of a test. The probabilityof a TypeII error

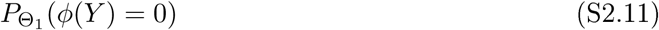

lacks a common denomination. Its complementary probability

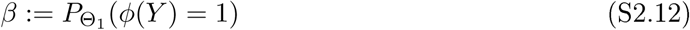

is referred to as the *power* of a test. In words, the power of a test is the probability of accepting the alternative hypothesis (rejecting the null hypothesis), if *θ* ∈ Θ_1_, i.e., if the alternative hypothesis is true. Note that basic introductions to test error probabilities often denote the probability of a Type II error by *β* ∈ [0, 1] and thus define power by 1 − *β*. For our current purposes, we prefer the definition of eq. (S2.12), because it keeps the notation concise and is more coherent with common notations of test quality functions.

##### SIGNIFICANCE LEVEL

It is important to distinguish between the size and the significance level of a test: a test is said to be of *significance level α′*∈ [0, 1], if its size *α* is smaller than or equal to *α′*, i.e., if

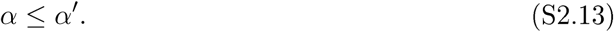

If for a test of significance level *α′* it holds that *α* < *α′*, the test is referred to as a *conservative test*. If for a test of significance level *α′* it holds that *α* = *α′*, the test is referred to as an *exact test*. Tests with an associated significance level *α′* for which *α* > *α′* are sometimes referred to as *liberal tests*. Note, however, that such tests are, strictly speaking, not of significance level *α′*.

##### THE TEST QUALITY FUNCTION

The size and the power of a test are summarized in the test’s quality function. For a test *ϕ*(*Y*), the *test quality function is* defined as

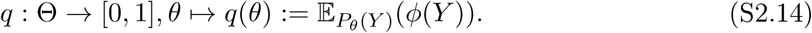

In words, the test quality function is a function of the probabilistic model parameter *θ* and assigns to each value of this parameter a value in the interval [0, 1]. This value is given by the expectation of the test *ϕ* under the probabilistic model *P*_*θ*_(*Y*). The definition of the test quality function is motivated by the value it assumes for *θ* ∈ Θ_0_ and *θ* ∈ Θ_1_: because the random variable *ϕ*(*Y*) only takes on values in {0, 1}, the expected value 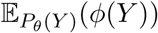 is identical to the probability of the event *ϕ*(*Y*) = 1 under *P*_*θ*_(*Y*). Thus, for *θ* ∈ Θ_0_, the test quality function returns the size of the test (eq. (S2.9)) and for *θ* ∈ Θ_1_, the test quality function returns the power of the test (eq. (S2.12)).

##### THE TEST POWER FUNCTION

For *θ* ∈ Θ_1_, the test quality function is also is referred to as the test’s *power function* and is denoted by

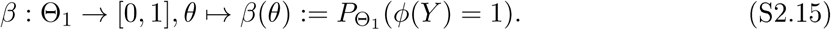

##### TEST CONSTRUCTION

In both applications and the theoretical development of statistical tests, the probability for a Type I error, i.e., the test size, is usually considered to be more important than the Type II error rate, i.e., the complement of the test’s power. In effect, when designing a test, the test’s size is usually fixed first, for example by deciding for a significance level such as *α′* = 0.05 and its associated critical value 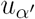 of the test statistic (cf. eq. (S2.8)). In a second step, different tests or different probabilistic models are then compared in their ability to minimize the probability of the test’s Type II error, i.e., maximize the test’s power. For example, the celebrated Neyman-Pearson lemma states that for tests of simple hypotheses, the likelihood ratio test achieves the highest power for a given significance level over all conceivable statistical tests (Neyman and Pearson, 1933). Inspired by current discussions about the power of tests in functional neuroimaging, in the current study we primarily target the sample size as a parameter of the probabilistic model to optimize different tests with respect to their Type II error rates given prefixed Type I error rates.

#### S.2.2 The multiple testing scenario

##### PROBABILISTIC MODEL

The notion of a multiple hypothesis test can be developed in analogy to the single test scenario. Like the single test scenario, the multiple testing scenario unfolds against the background of a parametric probabilistic model *P*_*θ*_(*Y*) that describes the probability distribution of a random entity *Y* which models observed data taking on values in ℝ^*n*^. The parameter *θ* of the model is assumed to take values in a parameter space Θ.

##### MULTIPLE TEST HYPOTHESES

In multiple testing scenarios comprising *m* ∈ ℕ tests, the parameter space is partitioned *m* times into disjoint subsets 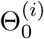 and 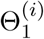, such that 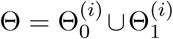 and 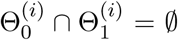 for *i* ∈ *I* := {1, …, m} and |*I* | = *m*. In analogy to the single test case, the statements

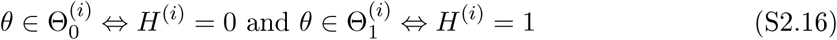

about the true, but unknown, value of the parameter *θ* are referred to as the *i*th *null* and *alternative hypothesis*, respectively. Collectively, the *m* null hypotheses and their associated alternative hypotheses are referred to as a *hypotheses family* and the set *I* is referred to as the *hypotheses index set*. In the following, we will be concerned with the following situations

- all null hypotheses of the hypotheses family are true and all alternative hypotheses are false,
- some null hypotheses of the hypotheses family are true and the remaining alternative hypotheses are true,
- all null hypotheses of the hypotheses family are false and all alternative hypotheses are true.

For convenience, we will refer to these scenarios as the *complete null hypothesis, the partial alternative hypothesis*, and the *complete alternative hypothesis*, respectively. The following notation is helpful to formally express the complete null hypothesis and complete alternative hypothesis scenarios, respectively:

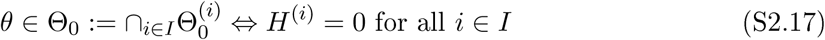

and

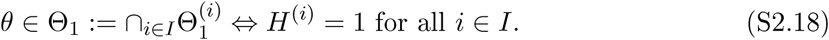

Note that despite the identical notation, the difference between the single test scenario null and alternative hypotheses (S2.1) and (S2.2), and the multiple testing scenario complete null and complete alternative hypotheses (S2.17) and (S2.18) should in general be clear from the context. As above, we will use the subscript notations 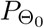 and 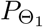 to indicate that the parameter *θ* of the probabilistic model *P*_*θ*_ is an element of the complete null or alternative hypotheses Θ_0_ or Θ_1_, respectively. In light of expressions (S2.17) and (S2.18), we denote the partial alternative hypothesis by

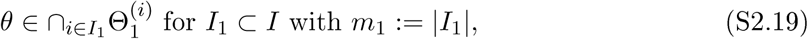

and refer to *I*_1_ as the *alternative hypotheses index set*. Given the binary nature of the *i*th null and alternative hypothesis, it follows immediately that in the case of (S2.19) it holds that

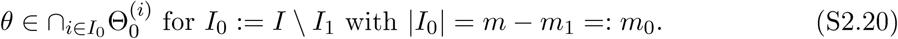

We refer to *I*_0_ as the *null hypotheses index set*. The ratio of the cardinality of the alternative hypotheses index set and the cardinality of the hypotheses index set will be denoted by

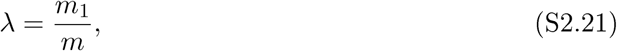

and will be referred to as the *alternative hypotheses ratio*. Note that *λ* = 0 corresponds to the complete null hypothesis, whereas *λ* = 1 corresponds to the complete alternative hypothesis. Finally, for *λ* ∈]0, 1[, we use the subscript notation 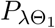 to indicate that the parameter *θ* of the probabilistic model *P*_*θ*_ is an element of a partial alternative hypothesis with alternative hypotheses ratio *λ*.

##### MULTIPLE TEST

For the multiple testing scenario, let

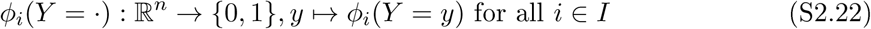

denote a test, such that *ϕ*_*i*_(*Y* = *y*) = 0 represents the act of accepting the *i*th null hypothesis and rejecting the *i*th alternative hypothesis, while *ϕ*_*i*_(*Y* = *y*) = 1 represents the act of rejecting the *i*th null hypotheses and accepting the *i*th alternative hypothesis. Then a multiple test is a mapping

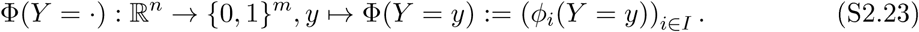

A multiple test can thus be conceived as an *m*-dimensional vector of single tests *ϕ*_*i*_(*Y* = ·), the probability distribution of which is governed by the parametric probabilistic model *P*_*θ*_(*Y*). As in the single test scenario, we will suppress the notational dependence of ϕ(*Y* = ·) on *y* and write Φ(*Y*) instead. Again, because the data *Y* is modelled as a random entity, the expression Φ(*Y*) should be read as a random vector. Similarly, as in the single test scenario we are only concerned with scenarios for which each constituent test *Φ*_*i*_(*Y*) of Φ(*Y*) is of the form

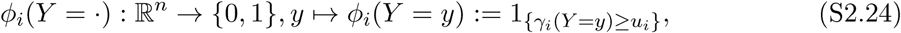

where

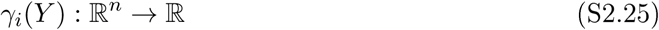

denotes the *i*th test statistic with *i*th rejection region

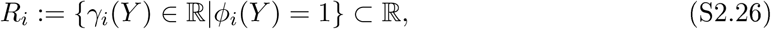

and *u*_*i*_ ∈ ℝ denotes the *i*th critical value. The multiple one-sided one-sample *T* -tests commonly performed for group-level fMRI analyses are a familiar example of the general multiple test structure described by eqs. (S2.23) - (S2.26): using voxel-specific sample means and sample standard deviations, the data *Y*, usually comprising voxel-wise participant-specific beta parameter estimate contrasts derived from first-level GLM analyses, is projected onto a set of *m T* -statistics. The values of these *m T* -statistics individually evaluated with respect to appropriately defined critical values, and for each of the *m* voxels, the null hypothesis of zero activation is either rejected or not.

**Table S2.**
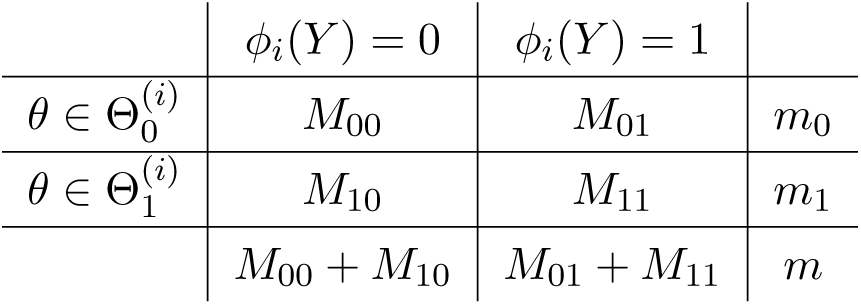
The multiple testing scenario. The numbers *m*_0_ and *m*_1_ of true null and alternative hypotheses 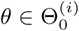 and 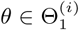, are assumed to be fixed and unknown. The outcome of the *i*th test *Φ*_*i*_(*Y*), and hence also the aggregate numbers of tests to assume either the value 0 or 1, *M*_*ij*_, *i* = 0, 1, *j* = 0, 1, as well as their sums, *M*_00_ + *M*_10_ and *M*_01_ + *M*_11_ are random entities, all of which are governed by the parametric probabilistic model *P*_*θ*_(*Y*) and the functional forms of the test statistics *γ*_*i*_, *i* = 1, …, *m*.

| | $\phi_i(Y) = 0$ | $\phi_i(Y) = 1$ | |
| --- | --- | --- | --- |
| $\theta \in \Theta_0^{(i)}$ | $M_{00}$ | $M_{01}$ | $m_0$ |
| $\theta \in \Theta_1^{(i)}$ | $M_{10}$ | $M_{11}$ | $m_1$ |
| | $M_{00} + M_{10}$ | $M_{01} + M_{11}$ | $m$ |

#### MULTIPLE TEST ERROR PROBABILITIES

The multiple testing scenario induces a variety of test error scenarios. While for the single test scenario there exist four possible constellations of true hypotheses and test outcomes (*θ* ∈ Θ_*j*_ and *Φ*(*Y*) = *k* for *j* = 0, 1 and *k* = 0, 1) there exist 4^*m*^ such constellations in the multiple testing scenario (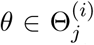 and *Φ*_*i*_(*Y*) = *k* for *i* = 1, …, *m, j* = 0, 1 and *k* = 0, 1). In other words, while a single test *Φ*(*Y*) may either result in either a Type I or a Type II error (or a correct result), a multiple test Φ(*Y*) may result in the simultaneous occurrence of Type I errors in some of its constituent single tests and Type II error in others of its constituents single tests (and correct results in the remaining single tests). This induces probabilities for the occurrence of a variety of test error scenarios and hence a variety of Type I and Type II error rates. As Type II error rates are complementary probabilities of correct rejections of null hypotheses, different Type II error rates correspond to different notions of power. In the following, we first review the most commonly considered Type I and Type II error rates in multiple testing scenarios. In later sections, we then consider the family-wise error rate, minimal and maximal power and their control and evaluation by means of maximum and minimum statistics in further detail.

The test error rates of multiple testing scenarios can be developed quantitatively as follows: as above, let *I*_0_ and *I*_1_ denote the null and alternative hypotheses index sets, respectively (cf. eqs. (S2.20) and (S2.19)). Note again that the binary single test scenario implies that *I* = *I*_0_ ∪ *I*_1_ and *I*_0_ ∩ *I*_1_ = ∅ and that it is assumed that the sets *I*_0_ and *I*_1_ and their respective cardinalities *m*_0_ and *m*_1_ are true, but unknown, entities. Based on the probabilistic binary outcome of each test constituent *Φ*_*i*_(*Y*), the following quantities are induced at an aggregate level:

- the number *M*_00_ of tests for which 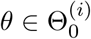 and *Φ*_*i*_(*Y*) = 0,
- the number *M*_01_ of tests for which 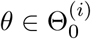 and *Φ*_*i*_(*Y*) = 1,
- the number *M*_10_ of tests for which 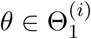 and *Φ*_*i*_(*Y*) = 0, and
- the number *M*_11_ of tests for which 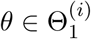 and *Φ*_*i*_(*Y*) = 1.

The situation is summarized in Table S.2. Note that the values *m, m*_0_ and *m*_1_ correspond to true, but unknown, quantities, the four quantities *M*_*jk*_, *j* = 0, 1, *k* = 0, 1 correspond to unobservable random variables, and the quantities *M*_00_ +*M*_10_ and *M*_01_ +*M*_11_, i.e., the total number of accepted and rejected null hypotheses, correspond to observable random variables. Commonly considered Type I error rates in this scenario are

- the *family-wise error rate*, defined as the probability for the event *M*_01_ ≥ 0, i.e., of one or more Type I errors,
- the *per-family error rate*, defined as the expectation of the unobservable random variable *M*_01_, i.e., the expected number of Type I errors,
- the *per-comparison error rate*, defined as the per-family error rate divided by the number of hypotheses *m*, and
- the *false-discovery rate*, defined as the expectation of the random variable *M*_01_*/*(*M*_01_ + *M*_11_) if *M*_01_ + *M*_11_ ≠ 0 and 0 if *M*_01_ + *M*_11_ = 0, i.e., the expected proportion of Type I errors among the rejected null hypotheses, or 0, if no hypotheses are rejected.

Notably, in contrast to the Type I error rate in the single test scenario (i.e., the size of a test), the Type I error rates in the multiple testing scenario refer to either probabilities (such as the family-wise error rate) or expectations of the counting random variables *M*_*ij*_, *i* = 0, 1, *j* = 0, 1. In a concrete multiple testing scenario, these probabilities and expectations have to be derived based on the nature of the probabilistic model and the definition of the multiple test.

As for the generalization of the notion of a Type I error to the multiple testing scenario, the multiple testing scenario induces a variety of Type II error rates and their respective complementary probabilities, i.e., power types. Commonly considered power types in the multiple testing scenario are

- *minimal power*, defined as the probability of the event *M*_11_ ≥ 1, i.e., of one or more correct rejections of the null hypothesis,
- *average power*, defined as the expectation of the random variable *M*_11_ divided by *m*_1_, i.e., the expected proportion of false null hypotheses that are rejected, and
- *maximal power*, defined as the probability of the event *M*_11_ = *m*_1_, i.e., of correctly rejecting all false null hypotheses.

##### MULTIPLE TEST CONSTRUCTION

As in the single test scenario, multiple tests are usually constructed to first and foremost control a chosen Type I error rate at a desired significance level *α ′*. In a second step, additional test construction measures may then be taken to achieve a desired level of a chosen power type. The random field theory-based fMRI inference framework has traditionally focussed on the family-wise error rate (FWER) as the target for Type I error rate control. In the following, we shall thus further elaborate on the definition of the FWER and establish how the distribution of the maximum statistic can be utilized for its control. Furthermore, we formally develop the notions of minimal and maximal power and their relation to maximum and minimum statistics, respectively.

##### MAXIMUM STATISTIC-BASED FWER CONTROL

As introduced above, the FWER of a multiple test is defined as the probability of one or more Type I errors. More formally, let Φ (*Y*) = (*Φ*_*i*_(*Y*))_*i*∈*I*_ denote a multiple test with with hypotheses index set *I* and null hypotheses index set *I*_0_ ⊆ *I, I*_0_ ≠ ∅. Then the FWER is defined as the probability

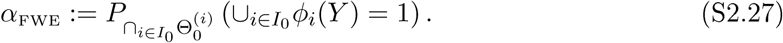

This expression is to be understood as follows: clearly, the FWER refers to the probability of events *Φ*_*i*_(*Y*) = 1 under the probabilistic model for the case that at least one null hypothesis holds true, i.e., *I*_0_ ≠ ∅. More specifically, the intersection subscript 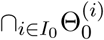 qualifies that the parameter of the probabilistic model is such, that all null hypotheses with indices in the set *I*_0_ ⊆ *I, I*_0_ ≠ ∅ hold. Complementary, the union statement 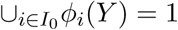 implies that the event 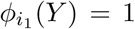 and/or the event 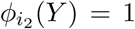, …, and/or the event 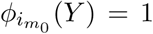 with *i*_*j*_ ∈ *I*_0_ for *j* = 1, 2, …, *m*_0_ occurs, i.e., that at least one, but possible more, events *Φ*_*i*_(*Y*) = 1 with *i* ∈ *I*_0_ occurs. This is equivalent to the probability of the event *M*_01_ ≥ 0 as considered above. In analogy to the significance level in the single test scenario, a multiple test Φ(*Y*) is then said to be of *family-wise significance level 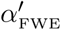*, if its FWER is equal to or smaller than 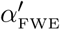, i.e., if

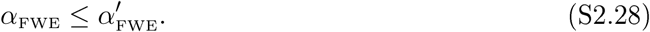

Equivalently, such a test is said to *control the FWER at level* 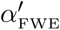. If for a test Φ(*Y*) it holds that 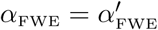, we say that Φ(*Y*) offers *exact control of the FWER at level* 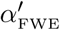 A general method to establish FWER control for a multiple test of the form (S2.23) - (S2.26) at a level 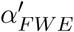 is afforded by consideration of the distribution of the *maximum test statistic*

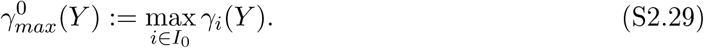

The method rests on identifying a common critical value 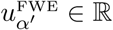 for all constituent tests *Φ* (*Y*) of the form (S2.24) that satisfies

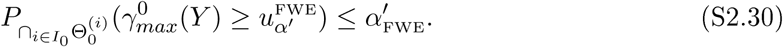

Intuitively, requirement (S2.30) states that the probability of the maximum of the multiple test’s test statistics to assume a value larger than or equal to the critical value 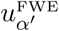 over the set of true null hypotheses is smaller or equal to the the desired FWER control level 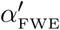 As shown below, from requirement (S2.30) it readily follows that

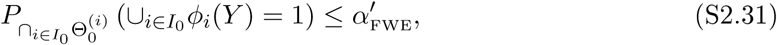

i.e., that the multiple test controls the FWER at level 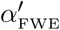. From an applied perspective, the maximum statistic-based FWER control approach entails that the distribution of the maximum statistic over the set of true null hypotheses needs to be evaluated based on the form of the probabilistic model and the resulting distributions of the component test statistics *γ*_*i*_(*Y*).

**Proof of eq. (S2.31)**

Let Φ(*Y*) denote a multiple test of the form eqs. (S2.23) - (S2.26), and define 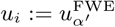 for all *i* ∈ *I*_0_ . Further, let the critical value 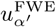 be such that with the definition of the maximum statistic 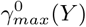 in eq. (S2.29) it holds that

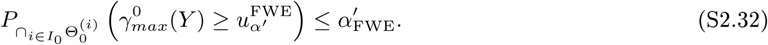

Then, with the definition of the FWER in eq. (S2.27), it follows that

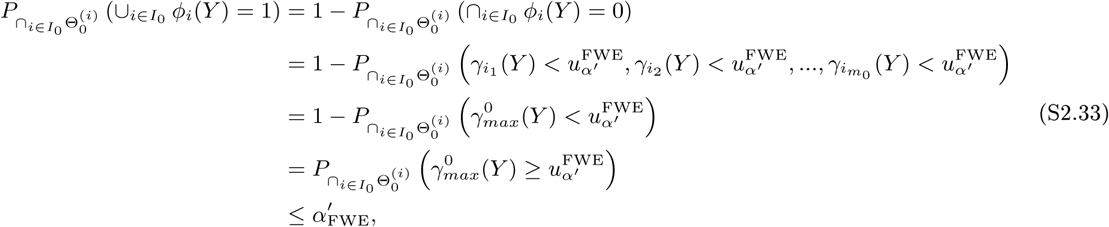

where on the right-hand side of the second equation *i*_*j*_ ∈ *I*_0_ for *j* = 1, 2, …, *m*_0_. In verbose form: the probability of the event that one or more of the component tests *Φ*_*i*_(*Y*), *i* ∈ *I*_0_ of the multiple test Φ(*Y*) evaluate to 1 over the set of true null hypotheses *I*_0_ is equal to the complementary probability of the event that all component tests evaluate to 0 over the set of true null hypotheses *I*_0_. Given the form of the multiple test Φ(*Y*), this probability in turn corresponds to the probability that all relevant component test statistics assume values smaller than the critical value 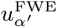 The latter event is identical to the event that the maximum statistic 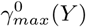 over the set of true null hypotheses is smaller than 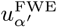 The complementary probability of this event then implies the validity of eq. (S2.31).

□

A note on the usage of the terms “uncorrected single test” and “corrected multiple testing” inference in the main text may be appropriate here: de-facto, FWER control in multiple testing is not based on some form of correction procedure that turns an “uncorrected *p*-value” into a “corrected *p*-value”, but the two *p*-values of uncorrected and corrected inference instead refer to different statistics. Because the notion of “correcting for the multiple testing problem” using “corrected *p*-values” is deeply engrained in the fMRI literature, however, we refrain from abandoning this terminology.

##### MAXIMUM STATISTIC-BASED MINIMAL POWER EVALUATION

Minimal power can be conceived of as the mirror analogue of the FWER. As defined above, minimal power is the probability for one or more correct rejections of the null hypothesis. In analogy to the FWER, minimal power of a multiple test Φ(*Y*) = (*Φ*_*i*_(*Y*))_*i*∈*I*_ with hypotheses index set *I* and alternative hypotheses index set *I*_1_ ⊆ *I, I*_1_ ≠ ∅ is formally given as

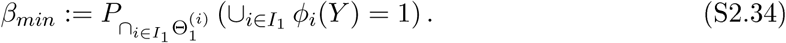

As for the formal expression of the FWER, the intersection statement 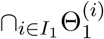 qualifies that the parameter of the probabilistic model is such that all alternative hypotheses with indices in the set *I*_1_ ⊆ *I, I*_1_ ≠ ∅ hold true, while the union statement 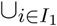 implies that the event 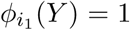 and/or the event 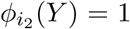, …, and/or the event 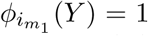 with *i*_*j*_ ∈ *I*_1_ for *j* = 1, 2, …, *m*_1_ occurs, i.e., that at least one, but possible more, events *Φ*_*i*_(*Y*) = 1 with *i* ∈ *I*_1_ occurs. This is equivalent to the event *M*_11_ ≥ 1 as considered above. Minimal power can be evaluated in a straight-forward fashion for multiple testing procedures that employ a common critical value and for which the distribution of the maximum statistic is known. Specifically, as shown below, given a test of the form (S2.23) - (S2.26), the definition of the maximum statistic

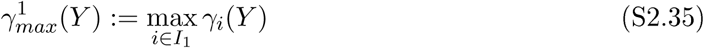

and a critical value *u* ∈ ℝ, it holds that

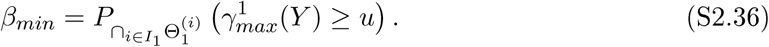

In the applied context of the current study, eq. (S2.36) implies that minimal power can be evaluated by considering the appropriate maximum statistic distributions of the random-field theory-based fMRI inference framework.

**Proof of eq. (S2.36)**

Let Φ be a multiple test of the form eqs.(S2.23) - (S2.26), let 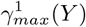 denote the maximum statistic as defined in eq. (S2.35), and let *u* ∈ ℝ denote an arbitrary critical value. Then eq. (S2.36) follows in analogy to the derivation of eq. (S2.31) as follows:

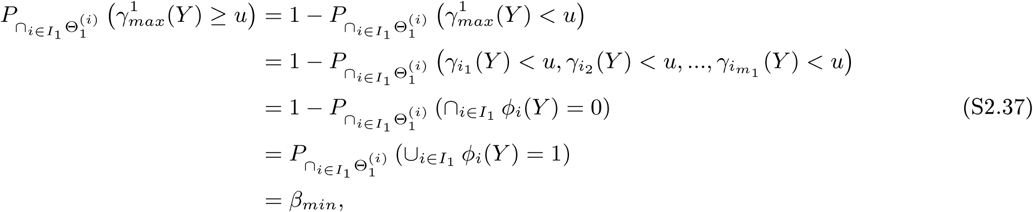

where on the right-hand side of the second equation *i*_*j*_ ∈ *I*_1_ for *j* = 1, 2, …, *m*_1_.

□

##### MINIMUM STATISTIC-BASED MAXIMAL POWER EVALUATION

In analogy to the formal definition of minimal power in eq. (S2.34), maximal power can be defined as

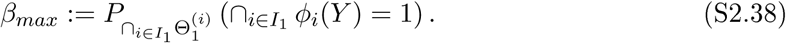

In analogy to the formal FWER and minimal power definitions, the intersection subscript 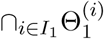 qualifies that the parameter of the probabilistic model is such, that all alterantive hypotheses with indices in the set *I*_1_ ⊆ *I, I*_1_ ≠ ∅ hold, while the intersection statement 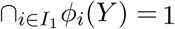 implies that the events 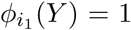 and the event 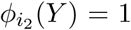, …, and the event 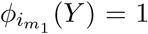 with *i*_*j*_ ∈ *I*_1_ for *j* = 1,2, …, *m*_1_ occur, i.e., that all events *Φ*_*i*_(*Y*) = 1 with *i* ∈ *I*_1_ occur. This is equivalent to the probability of the event *M*_11_ = *m*_1_ as considered above. Moreover, as shown below, given a test of the form (S2.23) - (S2.26), the definition of the *minimum* test statistic

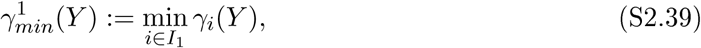

and a critical value *u* ∈ ℝ, it holds that

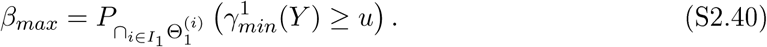

In the context of the current study, eq. (S2.40) implies that the evaluation of maximal power necessitates the availability of the minimum statistics distributions of the random-field theory-based fMRI inference framework under the appropriate alternative hypotheses scenarios.

**Proof of eq. (S2.40)**

Let Φ(*Y*) denote a multiple test of the form eqs.(S2.23) - (S2.26), let 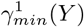 denote the minimum statistic as defined in eq. (S2.39), and let *u* ∈ ℝ denote an arbitrary critical value. Then eq. (S2.40) follows in analogy to the derivation of eq. (S2.36) as follows:

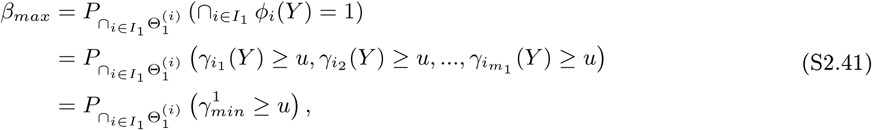

where on the right-hand side of the second equation *i*_*j*_ ∈ *I*_1_ for *j* = 1, 2, …, *m*_1_.

□

##### POWER FUNCTIONS

Based on the definitions of the partial alternative hypothesis ratio in eq. (S2.21), the definitions of minimal and maximal power in eqs. (S2.34) and (S2.38), respectively, and in analogy to the power function of the single test scenario (S2.15), we define the following *minimal* and *maximal power functions* of a given multiple test Φ(*Y*) with hypothesis index set *I, I*_0_ *⊂ I, I*_0_ ≠ ∅:

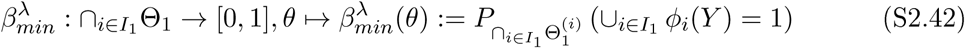

and

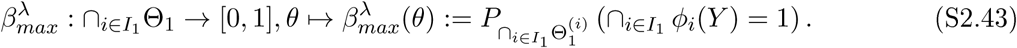

Note that the assumption *I*_0_ *⊂ I, I*_0_ ≠ ∅ implies that neither *I*_0_ nor *I*_1_ are empty sets and that hence *λ* ∈]0, 1[.

#### S.2.3 Positive predictive value functions

The concept of a positive predictive value (PPV) descends from a framework originally presented by Wacholder et al. (2004). Specifically, it arises in the context of probabilistic models, in which, in contrast to the classical frequentist test theory discussed thus far, both the test outcomes and the hypotheses states are modelled by random variables. In the following, we first consider the notion of a PPV in the context of the single test scenario discussed in Section S.2.1 and develop the notion of a PPV function. In a second step, we then consider the notion of a PPV in the multiple testing scenario of Section S.2.2 and define the ensuing minimal and maximal PPV functions.

##### THE SINGLE TEST SCENARIO

To establish the formal background of the PPV, we consider the parametric probabilistic model

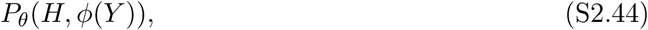

where the random variable *H* models the hypothesis state and the random variable *Φ*(*Y*) models the test state. As in Section S.2.1, *H* = 0 models the case that the null hypothesis is true and the alternative hypothesis is false, and *H* = 1 models the case that the null hypothesis is false and the alternative hypothesis is true (cf. eqs. (S2.1) and (S2.2)). Similarly, *Φ*(*Y*) = 0 represents the act of not rejecting the null hypothesis and *Φ*(*Y*) = 1 represents the act of rejecting the null hypothesis (cf. eq. (S2.3)). Note that the distribution of the data is considered only implicitly in the current probabilistic model, which is Justified as we again consider only deterministic test procedures. For the development of the PPV, the Joint distribution of *H* and *Φ*(*Y*) is constructed by (1) defining a parameterized marginal distribution for *H* and (2) employing the concepts of a single test’s size and power for the definition of the necessary conditional distributions. Specifically, the probability for the alternative hypothesis being true is parameterized by *π* ∈ [0, 1], inducing the marginal hypothesis prior distribution

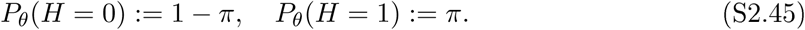

The required conditional distributions of *Φ*(*Y*) are then constructed by defining

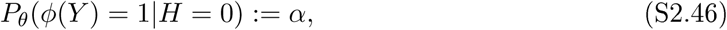

and

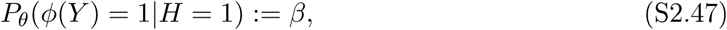

where *α* and *β* refer to the size and the power of the test *Φ*(*Y*) (cf. eqs. (S2.9)) and (S2.12), respectively. Based on the thus defined Joint distribution, the conditional probability of *H* to assume the value 1 given that *Φ*(*Y*) assumes the value 1 evaluates to

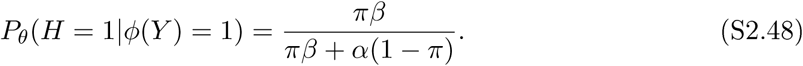

**Proof of eq. (S2.48)**

Eq. (S2.48) can be derived by (1) formulating the joint probability distribution *P*_*θ*_(*H, Φ*(*Y*)) based on the definition of the marginal distribution and conditional distributions *P* (*H*) and *P* (*Φ*(*Y*)*|H*) in eqs. (S2.45), (S2.46) and (S2.47), (2) evaluation of the marginal distribution *P*_*θ*_(*Φ*(*Y*)), and (3) evaluation of the ensuing conditional probability (S2.48).

(1) For the joint distribution, we have

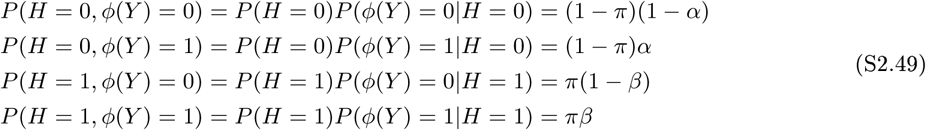

**Figure S.1.**
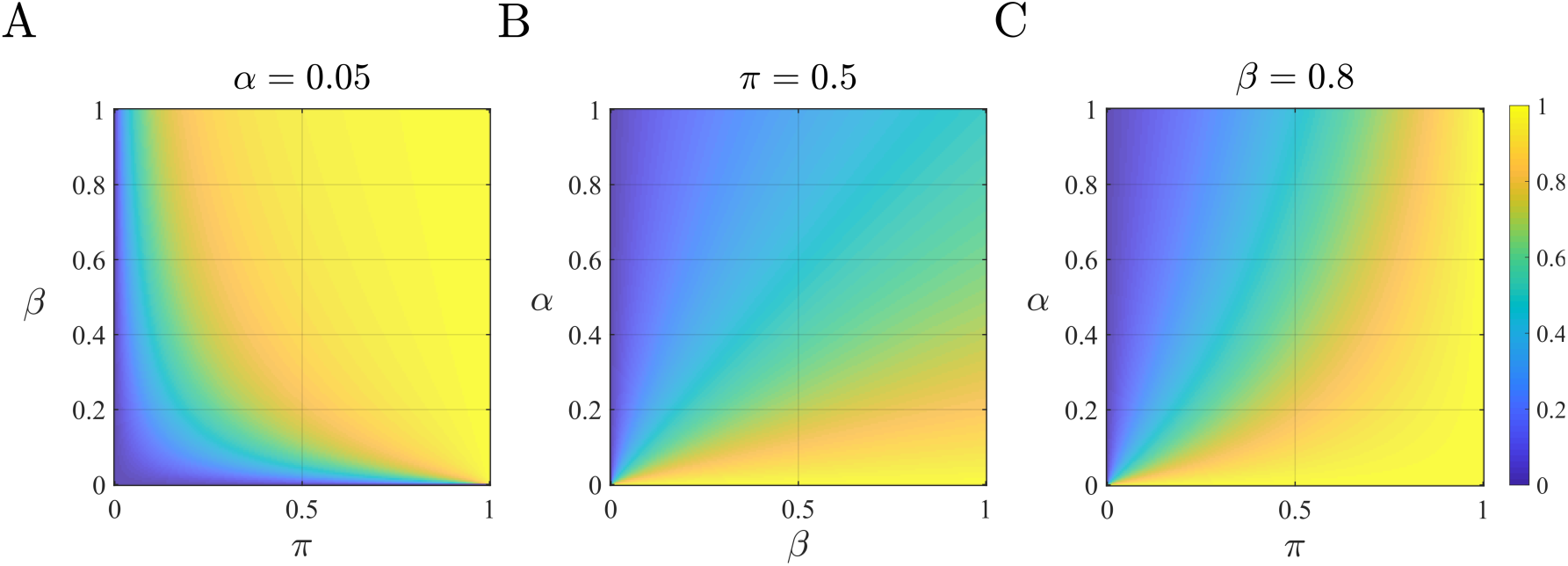
Positive predictive value. (**A**) Panel A visualizes the PPV as a function of the hypothesis prior probability *π* and the test power *β* for a fixed test size of *α* = 0.05. At fixed prior probability, an increase in power results in an increase of the PPV. For large prior probabilities, the effect of power on the PPV is negligible, while for low prior probabilities, the effect of power on the PPV is more pronounced. (B) Panel B visualizes the PPV for a uniform hypothesis prior probability distribution as a function of test power and test size. Optimal test properties of *α* = 0 and *β* = 1 result in an optimal PPV. For a test size of *α* = 0, optimal test power of *β* = 1 yields a PPV corresponding to the hypothesis prior probability *π* = 0.5. (C) Panel C visualizes the PPV for the commonly desired power of *β* = 0.8 as a function of the hypothesis prior probability and the test size. Note that the hypothesis prior probability dominates both test size and power. For implementational details, please see rftp_figure_S1.m.

(2) For the marginal distribution *P* (*Φ*), we thus have

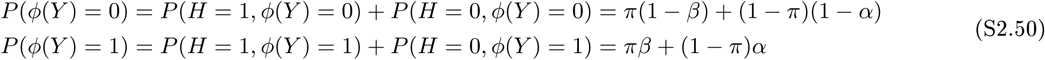

(3) Finally, for the conditional distribution of the alternative hypothesis being true (*H* = 1) given a positive test outcome *Φ*(*Y*) = 1, we have

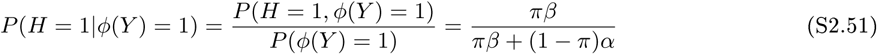

which completes the proof. □

For the probability *P*_*θ*_(*H* = 1*|Φ*(*Y*) = 1), Ioannidis (2005) coined the term positive predictive value. Intuitively, the PPV is thus the probability of the alternative hypothesis being true, given a positive test outcome. We visualize the dependency of the PPV on the hypothesis prior probability *π*, the test power *β*, and the test size *α* in Figure S.1. Fixing one of the three parameters of the PPV at a conventional level (*α* = 0.05, *π* = 0.5 and *β* = 0.8) demonstrates that the PPV of a test increases with the hypothesis prior probability and the test power, and decreases for increases in test size. Note that Ioannidis (2005) and Button et al. (2013) prefer the formulation of the PPV in terms of the *pre-study odds*

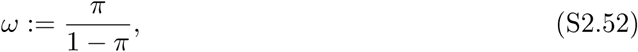

rather than hypothesis prior probability. In terms of the pre-study odds, the PPV can be re-expressed as

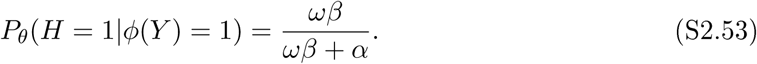

**Proof of eq. (S2.53)**

With the expression for the conditional probability of *H* = 1 given *Φ*(*Y*) = 1 and the definition of the pre-study odds of eq. (S2.52), we have

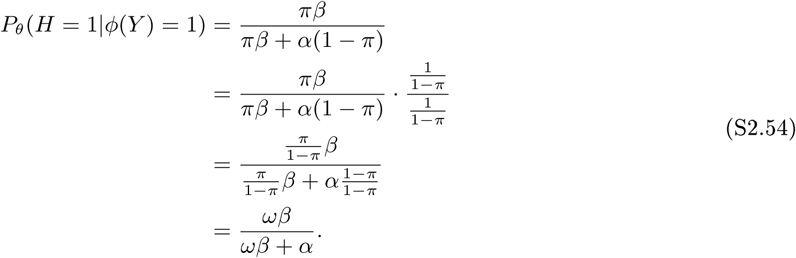

□

For a prefixed test size, the notions of a single test’s power function (cf. eq. (S2.15)) and the functional form of the PPV (cf. eq. (S2.48)) induce the positive predictive value function (PPV function)

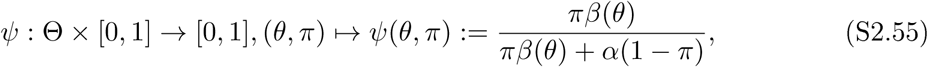

where *β*(*θ*) denotes the value of the test power function for *θ*. Note that the values of the PPV function depend on the prefixed test size *α*, the parameters of the probabilistic model by means of the single test power function *β*, and the hypothesis prior probability *π*.

##### THE MULTIPLE TESTING SCENARIO

To generalize the notion of a PPV function to the multiple testing scenario, we define the *minimal and maximal positive predictive value functions*

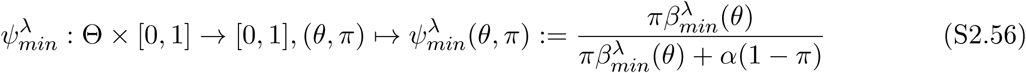

and

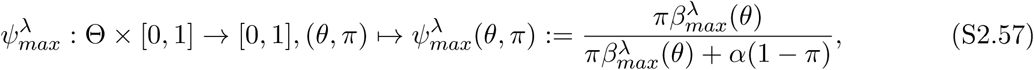

respectively, where *β*_*min*_ and *β*_*max*_ denote the minimal and maximal power functions as defined in (S2.42) and (S2.43). Note that in this scenario, the marginal hypothesis parameter *π* represents the prior probability of the partial alternative hypothesis scenario with partial alternative hypothesis parameter *λ* ∈]0, 1[.

#### S.2.4 Examples

To illustrate the theoretical concepts of Section S.2.1 to Section S.2.3 and as a conceptual reference point for the random field theory-based fMRI inference scenarios discussed in the main text, we next discuss two examples. The first example concerns a single test scenario, the second example concerns the extension of the first example to the multiple testing scenario. In both scenarios, we make repeated use of the probability density function of the Gaussian distribution, which we abbreviate by

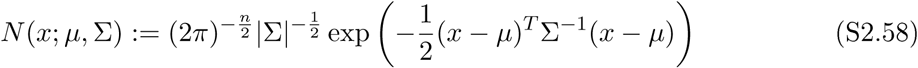

for expectation parameter *µ* ∈ ℝ^*n*^ and positive-definite covariance matrix parameter S ∈ ℝ^*n×n*^.

##### A single-observation z-test

###### PROBABILISTIC MODEL

As a first example, we consider a probabilistic model *P*_*θ*_(*Y*) that governs the distribution of a data random variable *Y* taking values in ℝ. For *µ* ∈ ℝ and *σ*^2^ > 0, the model is assumed to be defined in terms of the probability density function

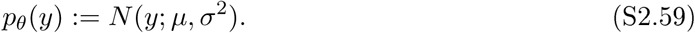

Intuitively, a single data point *Y* = *y* is thus assumed to have been sampled from a univariate Gaussian distribution of unknown expectation and known variance. For this model, we assume that the parameter space of interest is of the form Θ := ℝ_≥0_.

###### TEST HYPOTHESES1 STATISTIC1 AND DEFINITION

A single test scenario is then induced by defining the null and alternative hypotheses

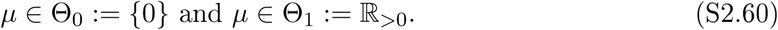

Furthermore, a test of the form (S2.8) can be constructed by defining the identity test statistic

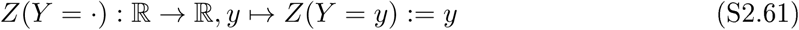

and the test

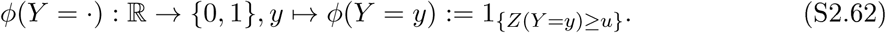

In words, the null hypothesis *µ* ∈ Θ_0_ is rejected, if the data realization is equal to or exceeds a given critical value *u* ∈ ℝ, otherwise it is not rejected.

###### DISTRIBUTIONS OF THE TEST STATISTIC

As discussed in Section S.2.1, to afford Type I error rate control and to evaluate the power of a thus controlled test, the distributions of the test statistic under the null and alternative hypotheses are central. The former distribution allows for identifying a critical value such that the size of the test maximally assumes a certain probability. The latter distribution allows for evaluating the probability of rejecting the null hypothesis under the scenario of the alternative hypothesis being true. In the current test scenario, the distribution of the test statistic under the null hypothesis *θ* ∈ Θ_0_, and hence also the probabilities for the equivalent events *Z*(*Y*) ∈ [*u, ∞* [and *Φ*(*Y*) = 1, can be readily inferred: because the test statistic conforms to the identity mapping, its distribution for *µ* ∈ Θ_0_ is given by the probability density function

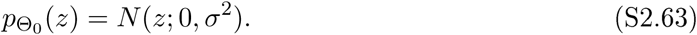

Likewise, the test statistic distribution for *θ* ∈ Θ_1_ and its associated events *Z*(*Y*) ∈ [*u, ∞* [and *Φ*(*Y*) = 1 is given by the probability density function

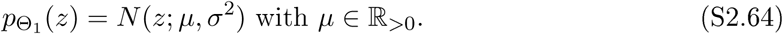

###### TYPE I ERROR RATE CONTROL

Given the form (S2.62) of the current test, *Φ*(*Y*) can be rendered an exact test of significance level *α ′* by choosing a critical value *u*_*α ′*_such that

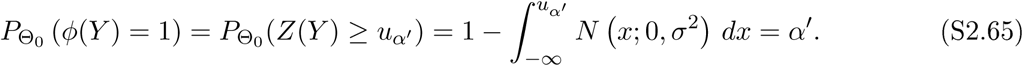

Note that the required integral corresponds to the cumulative density function of the univariate Gaussian distribution, for which well-known and widely implemented approximations exist. A numerical approach for the evaluation of *u*_*α ′*_based on the probability 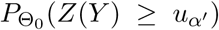 is discussed in Section S.6.

###### POWER AND POSITIVE PREDICTIVE VALUE FUNCTION

Given a critical value *u*_*α ′*_and the distribution of the test statistics under the alternative hypothesis scenario as specified by (S2.64), the probability of the event *Φ*(*Y*) = 1 evaluates to

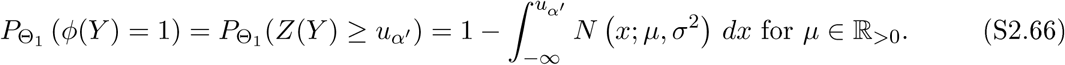

The power function of the test thus takes the form

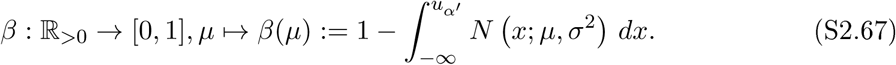

In applied settings, the parameterization of power functions in terms of the effect size measure *Cohen ’s d* is often preferred. For a univariate Gaussian distribution with expectation parameter *µ* ∈ ℝ and variance parameter *σ*^2^ > 0, Cohen’s *d* is defined as

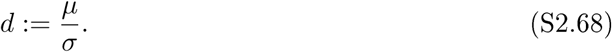

For the power function (S2.67), re-parameterization in terms of *d* results in

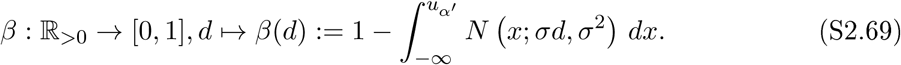

Finally, based on the form (S2.69) of the single-observation z-test power function and with the introduction of a prior hypothesis parameter *π*, the PPV function (cf. eq. (S2.55)) takes the form

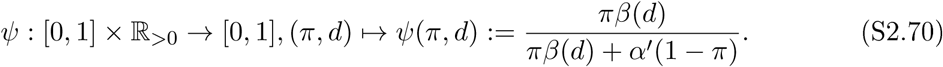

We visualize the single-observation z-test in Figure S.2. Figure S.2A visualizes the exceedance probability *P*_*θ*_(*Z* ≥ *z*) as a function of the statistic value *z* on the *x*-axis and the effect size *d* on the *y*-axis. In addition, the panel indicates the critical value *u*_*α ′*_= 1.645 for a significance level of *α ′* = 0.05 by a red line. The exceedance probabilities for *z* = *u*_*α ′*_as a function of the effect size *d* correspond to power function *β*, which is visualized in Figure S.2B. Finally, the PPV function ψ is visualized in Figure S.2C.

##### Multiple single-observation z-tests

###### PROBABILISTIC MODEL

We next consider the single-observation z-test in a multiple testing scenario. To this end, we assume a parametric probabilistic model *P*_*θ*_(*Y*) governing the distribution of a random vector *Y* = (*Y*_1_, …, *Y*_*m*_)*^T^* . Each component of *Y* is conceived as a univariate Gaussian random variable and the components are assumed to be distributed independently and with identical known variance *σ*^2^ > 0. A probability density function for the distribution of *Y* taking on values *y* ∈ ℝ^*m*^ is thus given by

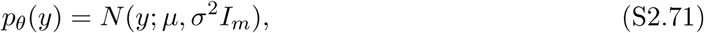

where *µ* ∈ ℝ^*m*^ and *I*_*m*_ denotes the *m × m* identity matrix. In other words, *Y* is distributed according to a multivariate Gaussian distribution with expectation parameter *µ* and spherical covariance matrix parameter *σ*^2^*I*_*m*_. The parameter space of the model is assumed to be given by 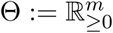 and concerns the expectation parameter *µ*.

**Figure S.2.**
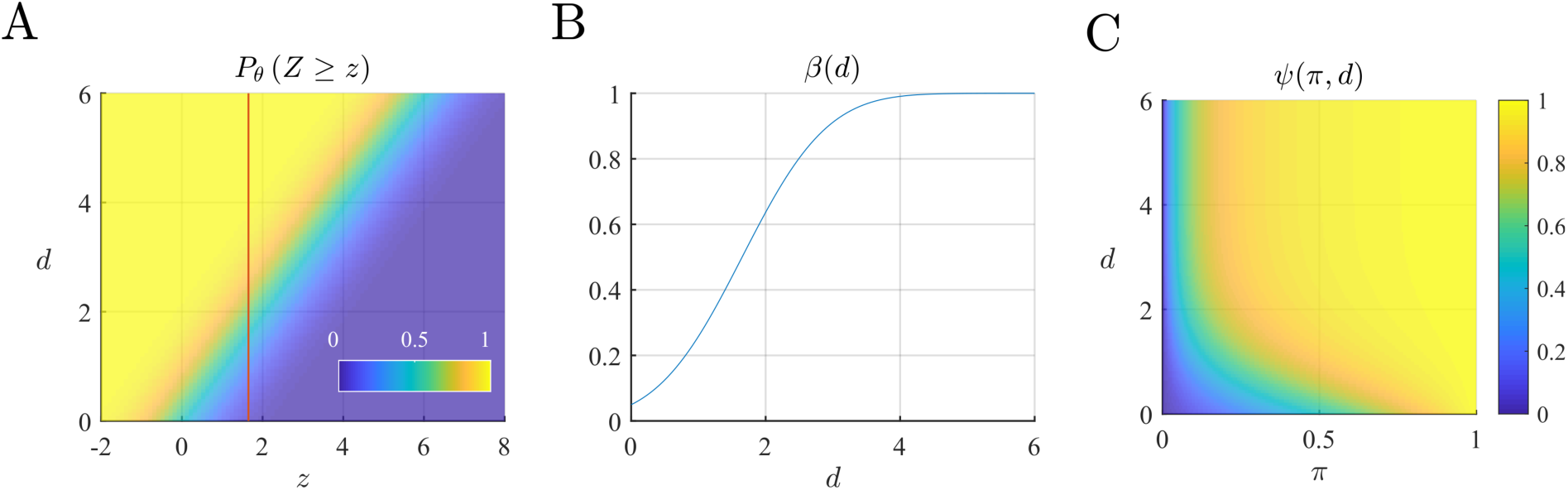
Exceedance probybility, power, and PPV functions for the single-observation z-test in the single test scenario. **(A)** The panel depicts the exceedance probability *P*_*θ*_ (*Z* ≥ *z*) as a function of the statistic value *z* and the effect size *d*. The red line indicates the critical value *u*_*α*_′ = 1.645 for a significance level of *α*′ = 0.05. (B) The power function of the single observation z-test. The values of the power function correspond to the the values of the EPFs for the critical value *u*_*α*_′ as depicted in Panel A. Note that for *d* = 0, the value of the power function corresponds to the size of the test. (C) The PPV function of the single-observation z-test. Note that for a hypothesis prior of *π* = 0, the PPV of the test does not exceed 0.5, while for a hypothesis prior of *π* = 1 the PPV of the test, is equal to one, regardless of the effect size. For implementational details, please see rftp_figure_S2.m.

###### TEST HYPOTHESES, STATISTICS, AND DEFINITION

For an index set *I* := {1, 2, …, *m*} and a value *θ* ∈ ℝ_>0_ we consider the family of hypotheses

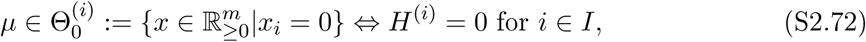

and

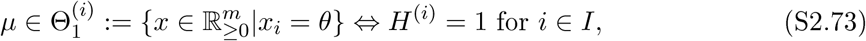

where *x*_*i*_ denotes the *i*th component of *x* ∈ ℝ^*m*^, *i* = 1, …, *m*. The *i*th null hypothesis thus states that the *i*th component of *µ* is zero and the remaining components of *µ* take on arbitrary values in ℝ_≥0_, while the *i*th alternative hypothesis states that the *i*th component of *µ* is equal to *θ* > 0 and the remaining components of *µ* take on arbitrary values in ℝ_≥0_. A multiple test of the form (S2.23) - (S2.26) can then be constructed by defining the test statistics

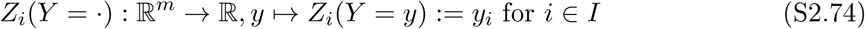

and defining the test

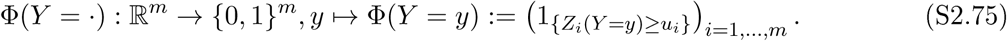

The *i*th test statistic thus corresponds to the proJection of *Y* onto its *i* coordinate. Note that for the current scenario the dimension of the data outcome space and the number of tested hypotheses are identical, but this does not necessarily have to be case.

###### DISTRIBUTIONS OF THE MAXIMUM STATISTIC

We next assume that we aim for the maximum statistic-based control of the FWER of the multiple test defined in eq. (S2.75). As discussed in Section S.2.2, this entails the evaluation of distribution the maximum statistic

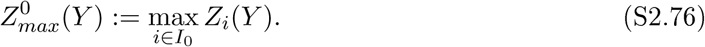

For the current example this distribution can be expressed in terms of the EPF

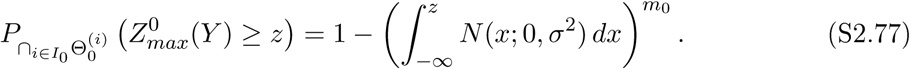

**Proof of eq. (S2.77)**

We have

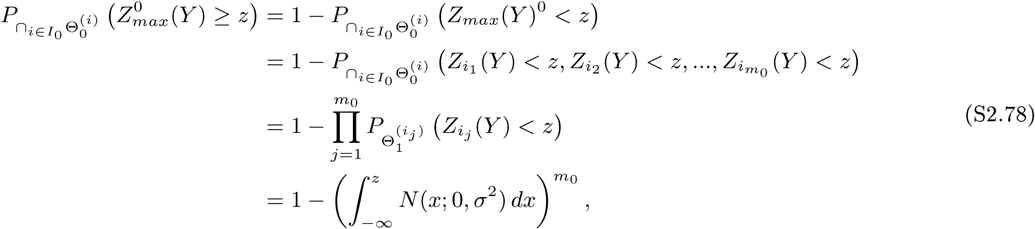

where *i*_*j*_ ∈ *I*_0_ for *j* = 1, 2, …, *m*_0_. The factorization of the joint distribution of the relevant test statistics implied by the third equation follows from the assumption of a spherical covariance matrix for the probabilistic model, which for the multivariate Gaussian distribution implies the independence of its component random variables. The fourth equation follows with the well-known form of the marginal distributions of the multivariate Gaussian distribution.□

Furthermore, we aim for the evaluation of minimal and maximal power functions and their associated PPV functions. As discussed in Section S.2.2, this entails the evaluation of the distributions of

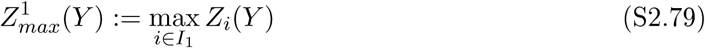

and

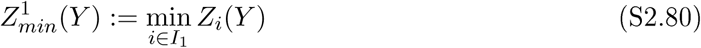

As shown below, these distributions can be expressed in terms of the EPFs

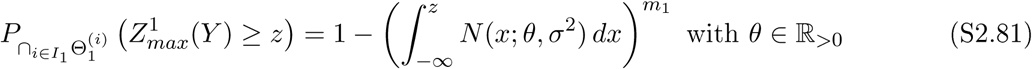

and

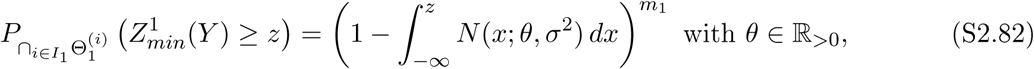

respectively.

**Proof of eqs. (S2.81) and** (S2.82)

The EPF (S2.81) follows as in the proof of (S2.77) by substitution of *I*_1_, 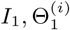 and *m*_1_ for *I*_0_, 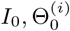 and *m*_0_, respectively. Similarly, the EPF (S2.82) follows from

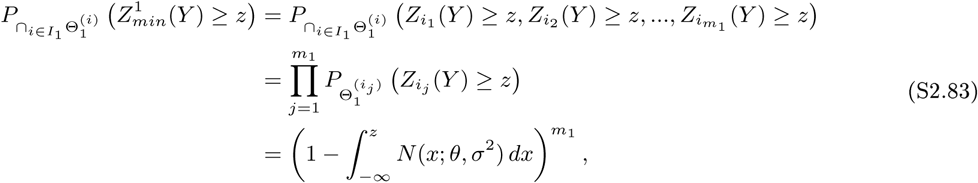

where *i*_*j*_ ∈ *I*_1_ for *j* = 1, 2, …, *m*_1_.□

**Figure S.3.**
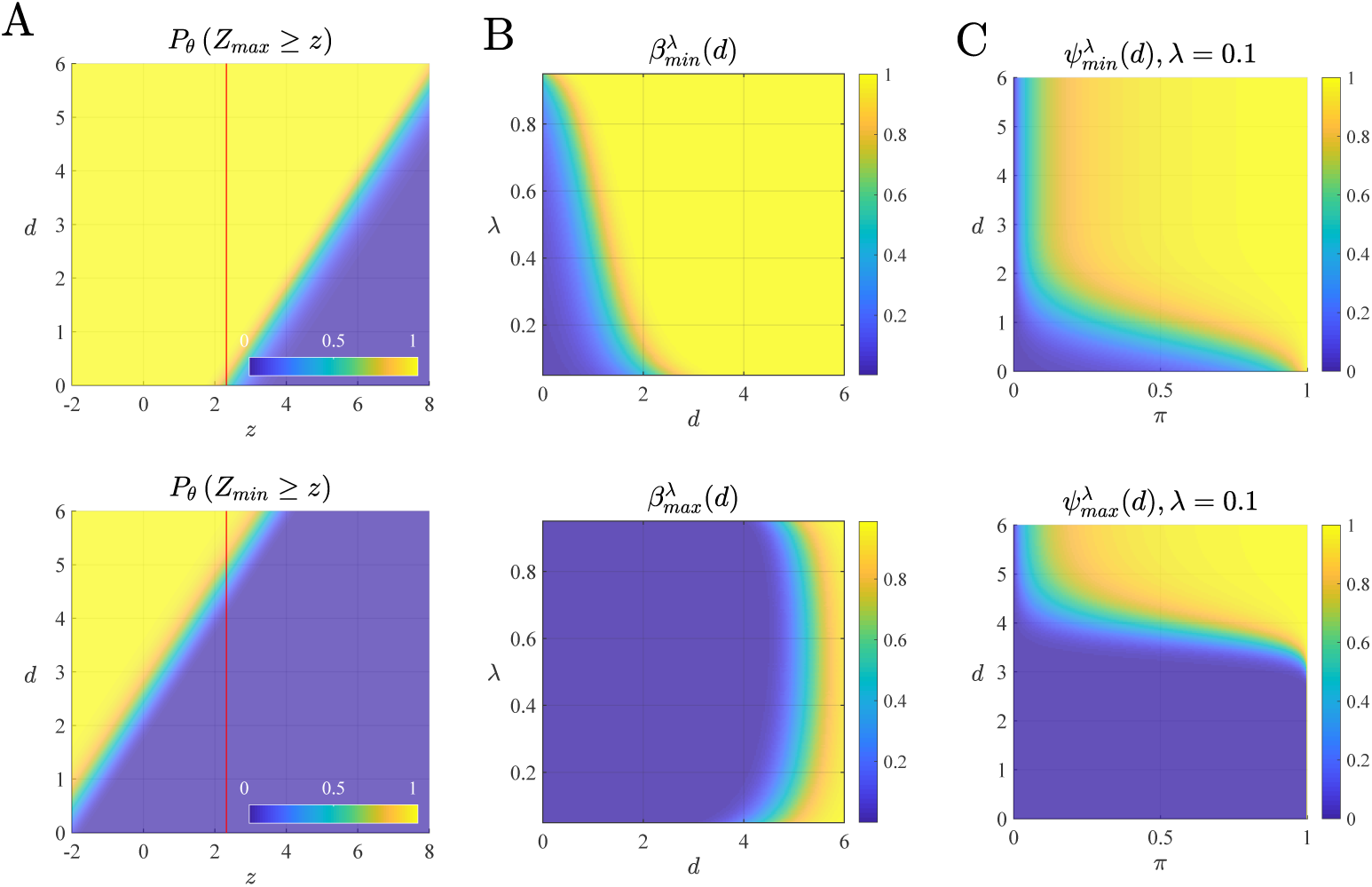
Exceedance probability, power, and PPV functions for the single observation z-test in the multiple testing scenario. (**A**) Maximum and minimum statistic EPFs for the multiple single-observation multiple z-test scenario with *m* := 100 hypotheses and variance parameter *σ*^2^ := 1. Note that when compared to the single test scenario, the critical value 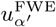 for a family-wise significance level of *α*_FEW_ = 0.05′ as indicated by the red lines assumes a value approximately twice as large. (**B**) Minimum and maximal power as functions of the partial alternative hypothesis parameter *λ* and the effect size parameter *d*. Note that to achieve similar levels, maximal power requires much larger effect sizes than minimal power. (**C**) Minimal and maximal PPVs as functions of the prior partial alternative hypothesis parameter and the effect size parameter *d* for a partial alternative hypothesis parameter of *λ* = 0.1. As in the single test scenario, extreme prior alternative hypothesis parameters render the PPV less dependent on the effect size than medium sized prior alternative hypothesis parameters. For implementational details, please see rftp_figure_S3.m.

###### TYPE I ERROR RATE CONTROL

As discussed in Section S.2.2, exact FWER control at significance level 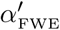 is afforded by identifying a common critical value 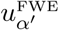 such that

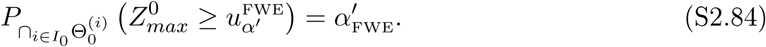

Given the form (S2.77) of the exceedance probability, the value of 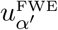 can be evaluated using the numerical approach discussed in Section S.6.

###### POWER AND POSITIVE PREDICTIVE VALUE FUNCTIONS

With the maximum statistic and minimum statistic dependencies of minimal and maximal power of eqs. (S2.36) and (S2.40), the parametric forms of the respective EPFs of eqs. (S2.81) and (S2.82), and the FWER controlling critical value (S2.84) it follows that the minimal and maximal power functions for the current example take the forms

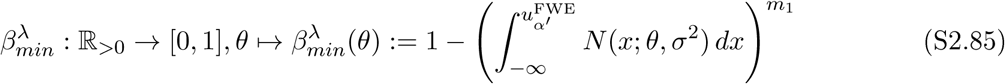

and

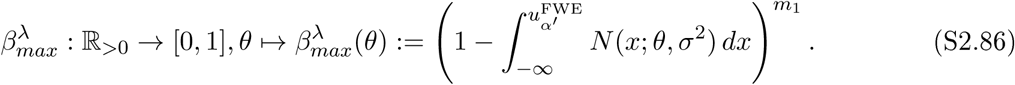

As for the single test scenario, reparameterization in terms of Cohen’s *d* results in the equivalent expressions

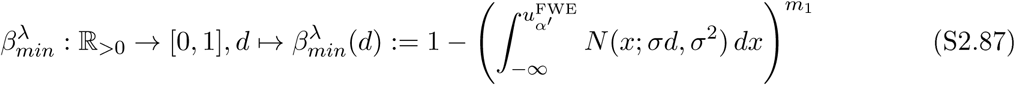

and

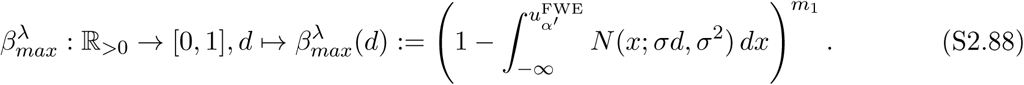

Finally, the introduction of a partial alternative hypothesis prior *π* induces the minimal and maximal PPV functions

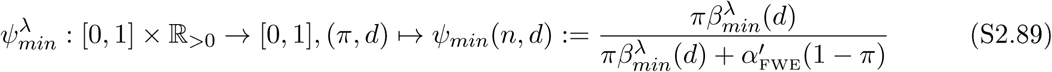

and

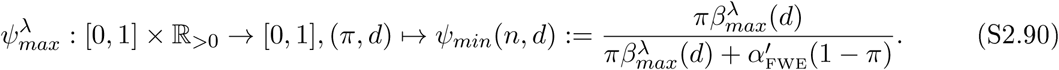

We visualize the multiple testing scenario of the single-observation z-test in Figure S.3 for the case of *m* := 100 simultaneously tested hypotheses. The upper and lower subpanels of Figure S.3A visualize the exceedance probabilities *P*_*θ*_ (*Z*_*max*_ ≥*z*) and *P*_*θ*_ (*Z*_*min*_ ≥ *z*). Note that in comparison with the *Z* statistic of the single test scenario in Figure S.2A, the maximum statistic exceedance probabilities are shifted to larger values of *z*, i.e., the maximum statistic *Z*_*max*_ has a higher probability to exceed a given *z* value than the *Z* statistic, and decays faster. Similarly, the minimum statistic exceedance probability mass is shifted to lower values of *z*, with the same decay as the maximum statistic. In addition, the subpanels of Figure S.3A indicate the critical value 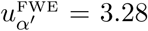 for a significance levelof *α*′^FWE^= 0.05 by a red line. Note that in comparison to the single test scenario, this critical value is approximately twice as large. The upper and lower panels of Figure S.3B visualize the ensuing minimal and maximal power function 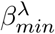 and 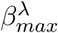, respectively, as a function of the effect size *d* and the partial alternative hypothesis parameter *λ* ∈]0, 1[. Note that for both power types, a high level of *λ* implies a small value of *m*_*0*_, which in turn results in a lower critical value 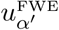, which for constant significance level and effect size, implies a higher value of the respective power function. This effect is particularly prominent in the case of maximal power, which for comparable power levels requires much higher effect size values when compared to minimal power, and exhibits a symmetry about a partial alternative hypothesis parameter of *λ* = 0.5. Finally, the upper and lower subpanels of Figure S.3C visualize the minimal and maximal PPV functions 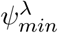 and 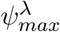 for *λ* = 0.1, respectively. Like for the single test scenario, the introduction of a partial alternative hypothesis scenario prior probability results in a modulation of the respective power functions, which for prior parameter values towards the boundaries *π* = 0 and *π* = 1 of the prior parameter space render the PPV function less dependent on effect sizes thanin the center of the prior parameter space around *π* = 0.5.

### S.3. Minimum statistics EPFs

#### Minimum voxel height statistic EPF

The approximation of the EPF of the minimum voxel height statistic is based on the assumption that for sufficiently high degrees of freedom the parametric expression of the expected Euler characteristic of a non-central *T* -field (cf. Hayasaka et al. (2007, eq. (4)), Worsley et al. (1996, eq. (3.1))) can serve both as an approximation for the probability of the maximum voxel height statistic to exceed a value *t* > 0, as well as as an approximation for the probability of the minimum voxel height statistic to fall below −*t*, i.e.,

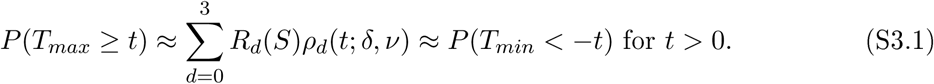

We then have for *t* > 0

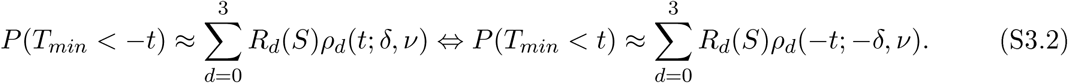

With the transformation *x* ≈ 1−exp(−*x*) for small *x* ∈ℝ that is used in the SPM implementation of RFT-based fMRI inference (cf. Friston et al. (1996, eq. (5)), Hayasaka et al. (2007, eq. (4)), Ostwald et al. (2018, eq. (110))), we then have

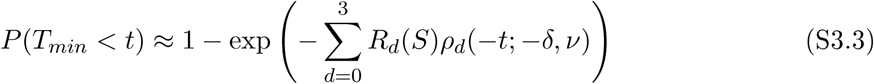

and hence

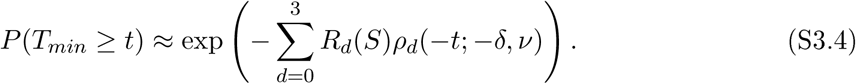

#### Minimum cluster extent statistic EPF

The approximation of the EPF of the minimum cluster extent statistic can be derived in analogy to the approximation of the EPF of the maximum cluster extent statistic (cf. Friston et al. (1994), Ostwald et al. (2018, Section 3.2)). To this end, let *M*_*u*_ denote the number of local maxima within an excursion set at level *u* of a non-central *T* -field, and let

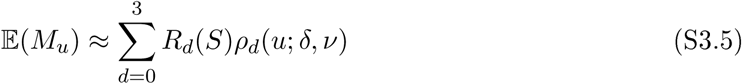

denote the expected Euler characteristic approximation of the number of local maxima within an excursion set at level *u* that also serves as the approximation to the expected number of clusters within an excursion set at level *u* under RFT-based fMRI inference. Further, let *C*_<*k,u*_ denote a random variable that models the number of clusters within an excursion set at level *u* that have an extent smaller than some constant *k*. As for its complement *C*_≥*k,u*_, which forms the basis for the approximation of the EPF of the maximum cluster extent statistic, we assume that

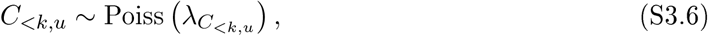

where

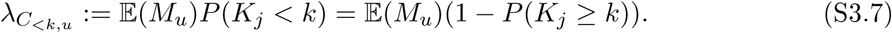

That is, like the random variable *C*_≥*k,u*_, the random variable *C*_<*k,u*_ is assumed to be distributed according to a Poisson distribution, the expectation parameter of which is given by the product of the expected number of clusters and, in contrast to *C*_≥*k,u*_, the probability of a cluster volume to take on a value smaller than some constant *k*. Next, let *K*_*j*_, *j* = 1, …, *c* denote the volumes of clusters *j* = 1, …, *c* within an excursion set at level *u*, and let *K*_*min*_ denote the minimum cluster extent statistic defined in eq. (7). Then, with the definition of the random variable *C*_<*k,u*_, we have for *k* ∈ ℝ

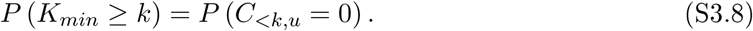

In words, the probability that the minimum of the cluster extent statistics *K*_*j*_ within an excursion set at level *u* is larger than or equal to *k* is identical to the probability that the number of clusters within the excursion set that have a volume smaller than *k* is zero. With the Poisson form of the distribution of *C*_<*k,u*_, it then follows that

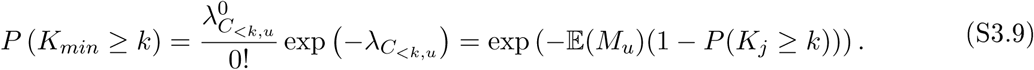

and with (S3.5), we obtain

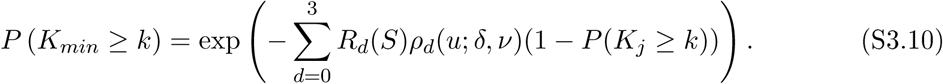

### S.4. EPFs visualizations

In this Section, we visualize EPFs of the six test statistics of eqs. (4) - (7) that underlie RFT-based fMRI inference and the power, PPV, and sample size calculations reported in the current work. These visualizations allow for readily relating the respective power functions to the distributional properties of the test statistics and in this way make the power functions discussed and documented in the main text accessible.

#### Single test scenario statistics

As discussed in Methods, the single test statistics of interest in RFT-based fMRI inference are the voxel height statistics *T*_*v*_ (cf. eq. (4)) and the cluster extent statistics *K*_*j*_ (cf. eq. (5)) with EPFs provided in eqs. (15) and (16), respectively. In Figure S.4A we visualize the test-relevant properties of the EPF of *T*_*v*_. Specifically, the upper panel of Figure S.4A depicts the critical value *t*_*α*_′ for exact tests with significance level *α*′ = 0.05 as a function of the sample size *n* implied by eq. (15) and evaluated using Algorithm 1. With increasing sample size, the critical value decreases, reflecting the lighter tail of the probability density function of Student’s *T* -distribution for high degrees of freedom. In the lower panel of Figure S.4A, we visualize the EPF of *T*_*v*_ as a function of effect size *d* and sample sizes *n* = 10, 15, …, 40. Here, each sample size-specific stack reflects a variation of the effect size *d* between 0.2 at the bottom of the stack and 0.8 at the top of the stack. Naturally, as the effect size increases, more probability mass is allocated to larger values of the test statistic outcome value *t*. In addition, the lower panel of Figure S.4A indicates the critical values for an exact test with significance level *α*′ = 0.05 at a given sample size by a red vertical bar. The value of the EPF at the location of this critical value for a given combination of sample and effect size corresponds to the power of the uncorrected voxel-level test as visualized in Figure 1A of the main text.

Similarly, the test-relevant aspects of the EPF of *K*_*j*_ are visualized in Figure S.4B for a cluster-defining threshold of *u* = 4.3. As for the voxel height statistic, the upper panel of Figure S.4B depicts the critical value *k*_*α*_′ measured in number of voxels for exact tests with significance level *α*′ = 0.05 as a function of the sample size *n* implied by eq.(16) and evaluated using Algorithm 1. As sample size increases, the critical cluster extent value decreases as expected. In the lower panel of Figure S.4B, we visualize the EPF of *K*_*j*_ as a function of effect size *d* and sample sizes *n* = 10, 15, …, 40. As in the lower panel of Figure S.4A, each sample size-specific stack reflects a variation of the effect size *d* between 0.2 (bottom) to 0.8 (top) and the critical values for an exact test with significance level *α*′ = 0.05 at a given sample size are indicated by red vertical bars. Like for voxel height statistic, the value of the EPF at the location of the critical values corresponds to the power of the uncorrected cluster-level test as visualized in Figure 1B of the main text.

#### Multiple testing statistics

As discussed in Methods, the multiple testing statistics of interest are the maximum and minimum voxel height statistics *T*_*max*_ and *T*_*min*_ (cf. eq. (6)) with EPFs provided in eqs. (17) and (16), respectively, as well as the maximum and minimum cluster extent statistics *K*_*min*_ and *K*_*max*_ (cf. eq. (7)), with EPFs provided in eqs. (19) and (20), respectively. As in the main text, we visualize test-relevant aspects of the EPFs in Figure S.5 based on the resel volumes *R*_0_(*S*) = 6, *R*_1_(*S*) = 33, *R*_2_(*S*) = 354, and *R*_3_(*S*) = 705, and, for the minimum and maximum cluster extent statistics, a cluster-defining threshold of *u* = 4.3.

The left upper panel of Figure S.5A depicts the critical values 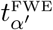 that afford FWER control at a significance level of 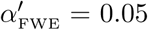 as a function of sample size *n* and the partial alternative hypothesis parameter *λ* ∈ [0, 1] for *d* = 0 and evaluated using Algorithm 1. For *λ* = 0 the scenario corresponds to the complete null hypothesis, and the critical values increase with decreasing sample size. Compared to the single test scenario, the critical values are five to ten times as large. Increasing *λ* and thus reducing the multiplicity of the multiple testing problem results in a decrease of the critical values, which accelerates as *λ* approaches 1. In the right upper panel of Figure S.5A, we visualize the associated exceedance probabilities that remain constant around 0.05. The medium panel of Figure S.5A visualizes the EPF of *T*_*max*_. As for *T*_*v*_ and *K*_*j*_, the EPFs are visualized as sample size-specific stacks for *n* = 10, 15, …, 40 and within each stack, the effect size *d* varies from 0.2 (bottom) to 0.8 (top). For all stacks, *λ* is set to 1, corresponding to the complete alternative hypothesis scenario. The red vertical bars reflect the sample size-specific critical values, and the values of the respective EPFs at the location of the vertical bars correspond to the minimal power values at the voxel level for *λ* = 1. Finally, the sample size-specific EPFs of *T*_*min*_ are visualized in the lowermost panel of Figure S.5A. Note that with respect to both the voxel height statistic (cf. Figure S.4A) and the maximum voxel height statistic (cf. Figure S.5A), considerably less probability mass is allocated to positive values of the statistic outcome value, reflecting the lower probability that the minimum of the voxel height statistics exceeds a given outcome value. As for *T*_*max*_, the red vertical bars indicate the FWER-controlling critical values, and the value that the minimum statistic EPF assumes at the location of these bars corresponds to the sample size- and effect size-specific minimal power of the FWER-controlled voxel-level test in the complete alternative hypothesis scenario. The identical way of portrayal is used in Figure S.5B for the test-relevant aspects of the EPFs of *K*_*min*_ and *K* _*max*_. Note that in comparison to the single test scenario, the critical values 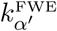 depicted in the upper left panel of Figure S.5B are up to three times as large.

**Figure S.4.**
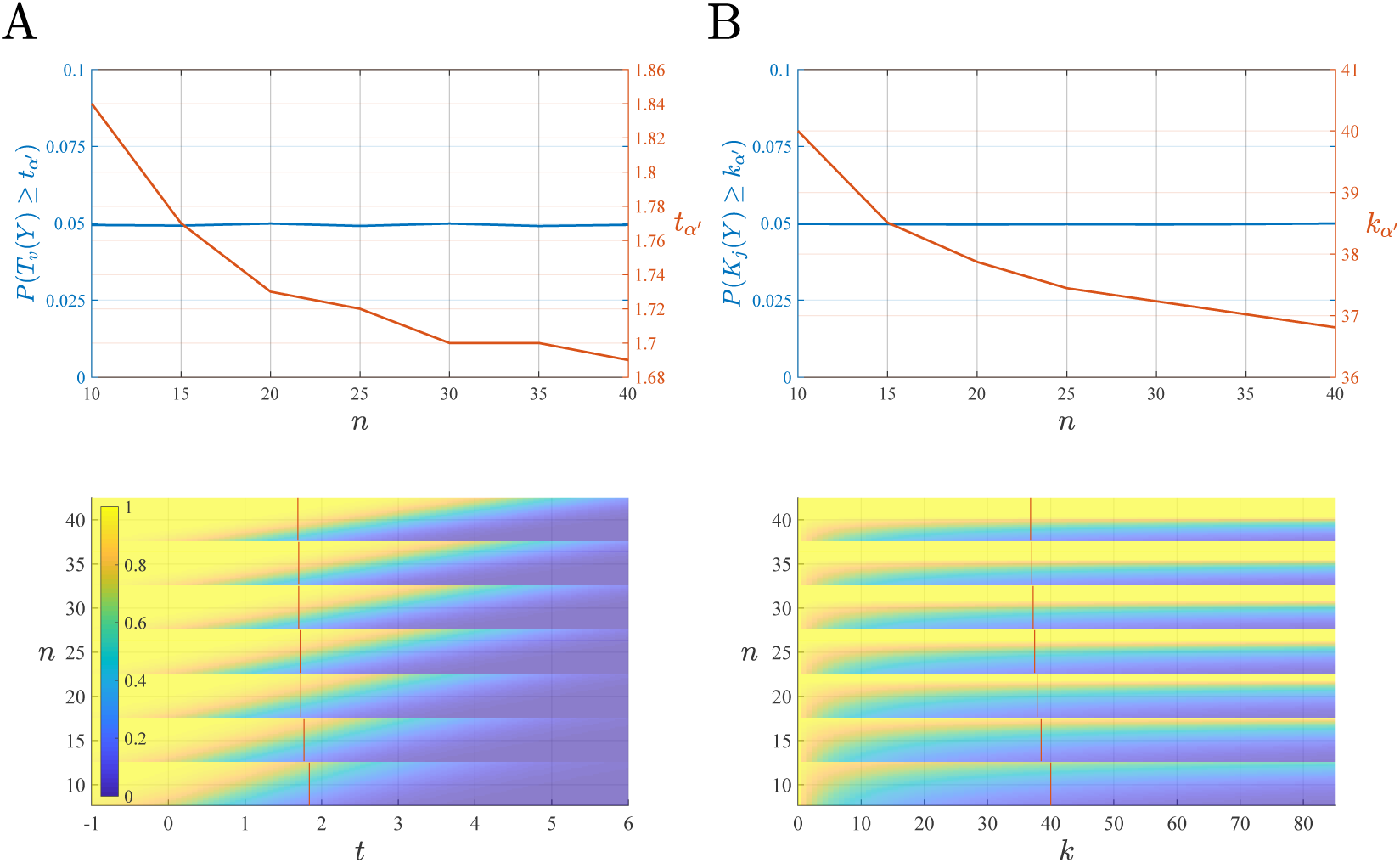
Critical values and exceedance probabilities for the single test voxel- and cluster-level statistics *Tv* and *K*_*j*_. The upper subpanels of panels (A) and (B) visualize the critical values *t*_*α*_′ and *k*_*α*_′ for exact single tests of significance level *α*′ = 0.05 as a function of sample size as red lines. These critical values were evaluated using Algorithm 1. The exceedance probabilities corresponding to these critical values are depicted as blue line and remain constant, as desired. The lower subpanels of panels (A) and (B) visualize the EPFs of eqs. (15) and (16) as a function of sample size and effect size *d*. Specifically, for each sample size-specific EPF stack, the effect size *d* varies between 0.2 at the bottom of the stack and 0.8 at the top of the stack. Additionally, the figure depicts the sample size-specific critical values for exact tests with significance level *α*′ = 0.05 as red vertical bars. The exceedance probabilities at the location of the red bars in the respective statistics outcome space corresponds to the effect and sample size-dependent power values visualized in Figure 1. Note that as discussed in the main text, neither EPF depends on resel volumes of the search space. For implementational details, please see rftp_figure_S4.m.

**Figure S.5.**
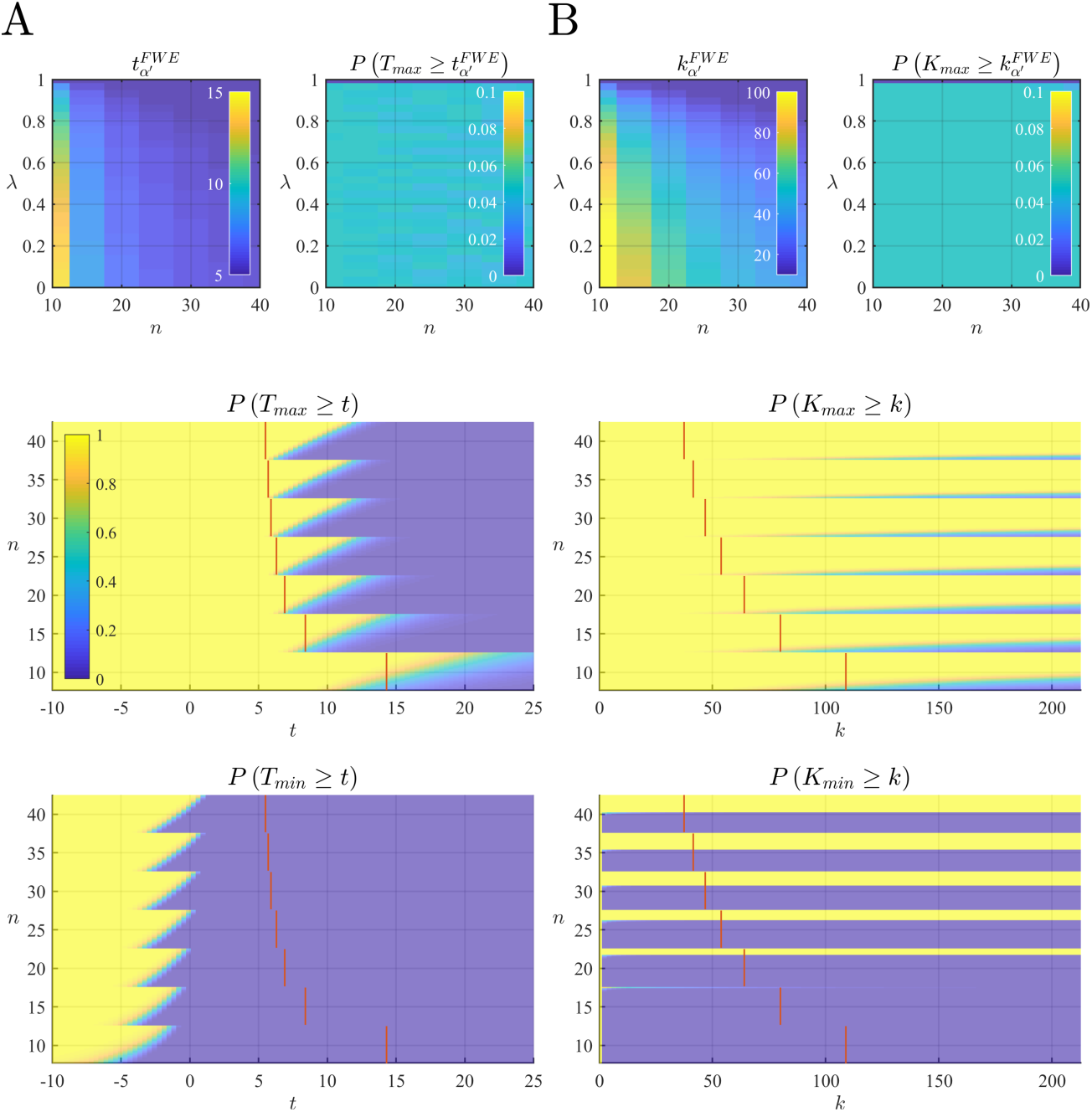
Critical values and EPFs for the multiple testing voxel- and cluster-level statistics. The left uppermost panels of (A) and (B) visualize the critical values 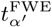 and 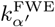 for FWER-controlled voxel- and cluster-level tests of significance level 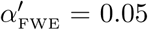 as a function of sample size *n* and partial alternative hypothesis parameter *λ* ∈ [0, 1], respectively. Their associated exceedance probabilities are visualized in the right upper panels of (A) and (B). The central panels visualize the EPFs of eqs. (17) and (19) as functions of sample size *n* and effect size *d*. Specifically, for each sample size-specific exceedance probability stack, the effect size *d* varies between 0.2 at the bottom of the stack and 0.8 at the top of the stack. In addition, each panel depicts the sample size-specific critical values for exact tests with significance level 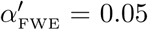 as red vertical bars. The color scale depicted for *P* (*T*_*max*_ ≥ *t*) applies to all four lower subpanels of the figure. The lowermost subpanels of (A) and (B) visualize the EPFs of eqs. (18) and (20) as functions of sample size *n* and effect size *d* as for their maximum counterparts. For implementational details, please see rftp_figure_S5.m.

### S.5. Exemplary data set

The exemplary fMRI data set is part of a perceptual decision making simultaneous EEG/fMRI data set that has been previously documented and made generally accessible in the standardized BIDS format (Ostwald et al., 2012; Georgie et al., 2018). In the following, we briefly sketch the experimental procedures and fMRI data analyses that form the basis for the statistical parametric map depicted in Figure 4.

#### Experimental procedure

Participants performed a visual perceptual decision task in a 2 × 2 factorial within-participant design with experimental factors “stimulus coherence” (with levels “low” and “high”) and “spatial prioritization” (with level “yes” and “no”). On each trial, a visual stimulus depicting either a face or a car was presented in one visual hemifield. Individual stimuli were presented for 200 ms and the participant was asked to indicate via a button press whether the stimulus depicted a face or a car. For the button presses, participants used their right index and middle finger for the two stimulus categories, and the mapping from stimulus category to response button was counterbalanced across participants. The informativeness of the visual stimulus was manipulated by altering the phase coherence of its spatial frequency spectrum resulting in low and high stimulus coherence trials. On half of the trials, a cueing arrow shown continuously for 1 s prior to the stimulus indicated in which visual hemifield the stimulus would be presented. Participants were asked to allocate their spatial attention to the respective visual hemifield, while maintaining steady central fixation (spatial prioritization condition). On the other half of the trials, the two-headed cuing arrow was uninformative and the stimulus was presented randomly in either visual hemifield (no spatial prioritization condition). Face and car stimuli were equally distributed across the four experimental conditions. The stimulus presentation order was randomized. Participants were asked to respond as quickly and as accurately as possible with an emphasis on responding as quickly as possible and to maintain stable fixation on the central fixation cross throughout the experiment. For fMRI data acquisition, data from 90 trials for each of the four conditions (half of them face stimuli) were recorded with an inter-trial interval discretely randomized between 10 and 12 s. The 90 trials per condition were split into five experimental runs, each lasting approximately 14 minutes.

#### fMRI data acquisition and analysis

fMRI data was acquired simultaneously with EEG at the Birmingham University Imaging Centre using a 3T Philips Achieva MRI scanner. T2*-weighted functional data were collected with an eight-channel phased-array SENSE head coil. EPI data (gradient echo-pulse sequence) were acquired from 32 slices (3×3×4 mm resolution, TR 2,000 ms, TE 35 ms, SENSE factor 2, flip angle 80 deg). Slices were oriented parallel to the AC-PC axis of the participant’s brain and positioned to cover the entire brain space. A mass-univariate summary-statistics GLM analysis was performed to assess condition-induced effects at the group-level. SPM12 (V6906) was used for both fMRI data preprocessing and statistical modelling. Prior to GLM parameter estimation at the participant-level, fMRI data were motion-corrected by realigning EPI volumes to the first volume of the first run of a given participant, normalized to MNI spaced using the SPM MNI-EPI template, re-interpolated to 2 mm isotropic voxel size, and smoothed using an 8 mm isotropic Gaussian kernel. The first-level GLM design matrix for each participant was then specified in run-wise, block-diagonal form. Here, each block comprised the four condition-specific stimulus onset functions, convolved with the canonical haemodynamic response function, in the column-wise order: high stimulus coherence/spatial prioritization, high stimulus coherence/no spatial prioritization, low stimulus coherence/spatial prioritization, low stimulus coherence/no spatial prioritization. Per SPM defaults, the design matrices additionally comprised a constant run offset and a cosine basis function set implementing a temporal high-pass filter with a cut-off of frequency 1/128 Hz. High-frequency residual error correlations were accounted for by SPM’s default of approximating a first-order autoregressive process with white noise using parameterized covariance basis functions. GLM beta and covariance component parameters were then estimated using SPM’s restricted maximum likelihood estimation scheme. Finally, ten participant-specific COPE images were evaluated for the “high stimulus coherence > low stimulus coherence” contrast weight vector (1, 1, −1, −1) replicated over sessions and padded with zeros for regressors of no interest. The resulting COPE images are available from the ‘Contrast Images’ folder of the accompanying OSF project. Finally, the COPE images were evaluated at the group-level using voxel-wise one-sample *T* -tests, implemented in rftp_figure_4.m, the resulting statistical parametric map of which is available from the ‘One Sample T Test’ folder in the accompanying OSF project.

### S.6. Algorithms

#### Critical value algorithm for a desired significance level

For a given test statistic *γ*(*Y*) and desired significance level *α*′, we use Algorithm 1 to numerically evaluate the required critical value *c*_*α*_′ for a test that controls the Type I error rate at significance level *α*′.

##### Algorithm 1 Critical value evaluation for a desired significance level

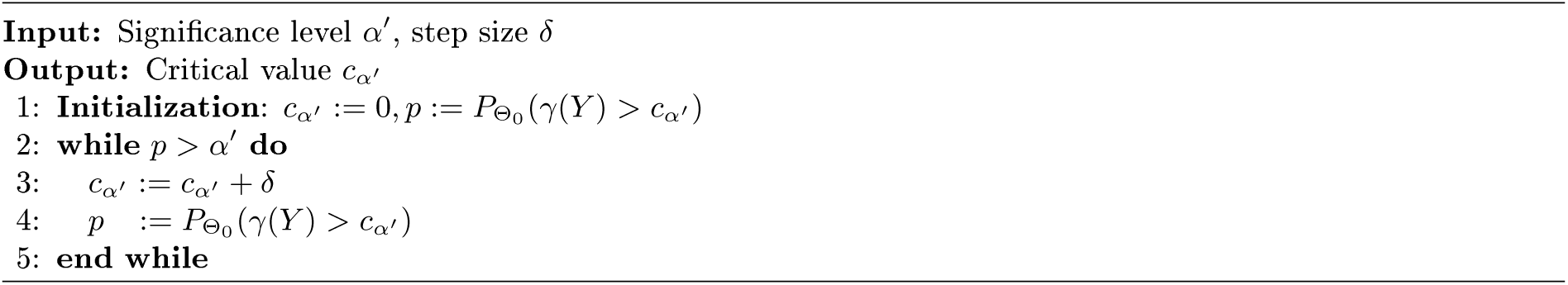

Algorithm A1 is implemented in the function u_fun.m.

#### Necessary sample size algorithm for a desired power level

For a given test statistic *γ*(*Y*), desired significance level *α*′, effect size *d* and, in the case of a multiple testing scenario, partial alternative hypothesis parameter *λ*, we use Algorithm 2 to numerically evaluate the required minimal sample size to achieve a desired power level *β*.

##### Algorithm 2 Necessary sample size evaluation for a desired power level

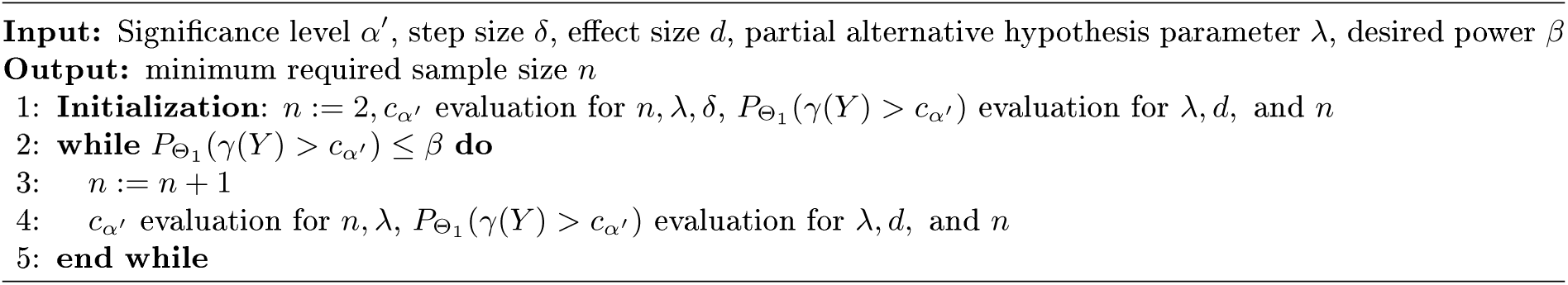

Algorithm A2 is implemented in the function n_fun.m.

#### Necessary sample size algorithm for a desired PPV level

For a given test statistic *γ*(*Y*), desired significance level *α*′, effect size *d*, prior hypothesis parameter *π* and, in the case of a multiple testing scenario, partial alternative hypothesis parameter *λ*, we use Algorithm 3 to numerically evaluate the required minimal sample size to achieve a desired PPV level *ψ*.

##### Algorithm 3 Necessary sample size evaluation for a desired PPV level

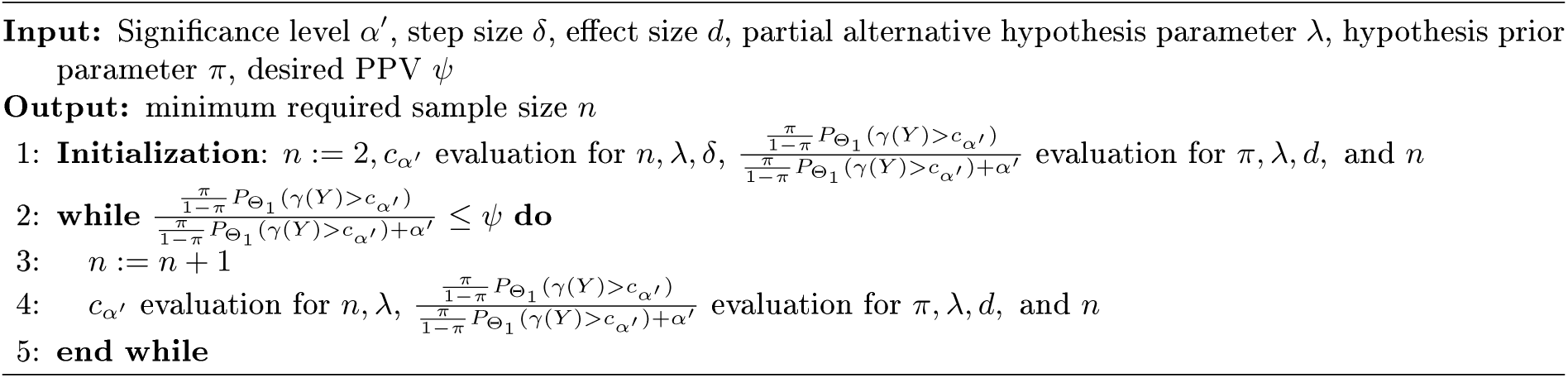

Algorithm A3 is implemented in the function n_fun.m.

